# Heart-on-a-chip model of immune-induced cardiac dysfunction reveals the role of free mitochondrial DNA and therapeutic effects of endothelial exosomes

**DOI:** 10.1101/2023.08.09.552495

**Authors:** Rick Xing Ze Lu, Naimeh Rafatian, Yimu Zhao, Karl T. Wagner, Erika L. Beroncal, Bo Li, Carol Lee, Jingan Chen, Eryn Churcher, Daniel Vosoughi, Ying Wang, Andrew Baker, Uriel Trahtemberg, Bowen Li, Agostino Pierro, Ana C. Andreazza, Claudia C. dos Santos, Milica Radisic

## Abstract

Cardiovascular disease continues to take more human lives than all cancer combined, prompting the need for improved research models and treatment options. Despite a significant progress in development of mature heart-on-a-chip models of fibrosis and cardiomyopathies starting from induced pluripotent stem cells (iPSCs), human cell-based models of myocardial inflammation are lacking. Here, we bioengineered a vascularized heart-on-a-chip system with circulating immune cells to model SARS-CoV-2-induced acute myocarditis. Briefly, we observed hallmarks of COVID-19-induced myocardial inflammation in the heart-on-a-chip model, as the presence of immune cells augmented the expression levels of proinflammatory cytokines, triggered progressive impairment of contractile function and altered intracellular calcium transient activities. An elevation of circulating cell-free mitochondrial DNA (ccf-mtDNA) was measured first in the *in vitro* heart-on-a-chip model and then validated in COVID-19 patients with low left ventricular ejection fraction (LVEF), demonstrating that mitochondrial damage is an important pathophysiological hallmark of inflammation induced cardiac dysfunction. Leveraging this platform in the context of SARS-CoV-2 induced myocardial inflammation, we established that administration of human umbilical vein-derived EVs effectively rescued the contractile deficit, normalized intracellular calcium handling, elevated the contraction force and reduced the ccf-mtDNA and chemokine release via TLR-NF-kB signaling axis.

## Introduction

Whereas numerous studies report bioengineering models of fibrosis and cardiomyopathies^1–3^, inflammatory conditions of the heart such as myocarditis are largely understudied. Most of the current mechanistic understanding of myocarditis comes from experiments in differentially susceptible rodent strains, which inevitably raises questions regarding the applicability of rodent-derived cardiac functional and immune system responses to human cardiovascular physiology. The absence of definitive therapeutic interventions for inflammatory heart conditions, such as myocarditis, is exacerbated by the lack of reliable biomarkers, difficulties in reaching confirmatory diagnosis and paucity in predictive human models for therapeutics testing. These factors collectively underscore the imperative nature of our work here. The successful modeling of heart inflammation hinges on the amalgamation of key advancements including 1) high-fidelity induced pluripotent stem cell (iPSC)-derived functional cardiac tissues, 2) establishment of a perfusable vasculature, 3) inclusion of immune cells, and 4) the ability to gather functional readouts such as contractility, calcium handling and electrical excitability in a single system.

While the clinical occurrence of acute myocarditis following SARS-CoV-2 infection is well-documented, the underlying biological mechanisms remain elusive. It was initially hypothesized that the internalization of viral RNA into the cardiomyocytes directly via the angiotensin-converting enzyme 2 (ACE2)^4, 5^ causes heart damage by impairing electromechanical functions^6–8^. As the COVID19 pandemic progressed, it became increasingly appreciated that systemic inflammation secondary to SARS-CoV-2 infection leads to cardiac complications, further motivating the development of human cell based vascularized disease models.

Complex conditions, such as viral induced myocardial inflammation, require new therapeutics. We focused on extracellular vesicles (EVs) as these subcellular entities, with an average diameter ranging between 80-200 nm, are pivotal mediators of intercellular communication. They achieve this by encapsulating and transferring selective biomolecules - miRNA, mRNA, proteins, and metabolites – from their parent cell to recipient cells, all without necessitating direct cell-to-cell contact^9^. Additionally, EVs are intrinsically biocompatible, biodegradable, and exhibit minimal cytotoxicity. Furthermore, EVs derived from human umbilical endothelial cells (HUVECs) have previously demonstrated promising regenerative, anti-inflammatory, and cardioprotective properties in pre-clinical settings^10–12^, hence their selection for inflammation management in our model.

Here, we used an organ-on-a-chip system, termed Integrated Vasculature for Assessing Dynamic Events (InVADE), which integrates iPSC-derived cardiomyocytes, an endothelialized micro-vessel structure termed Angiotube and peripheral blood mononuclear cells (PBMCs) to capture the complex cascade of events driving immune cell-induced cardiac dysfunction. We used SARS-CoV-2 as an example viral challenge, inducing acute myocarditis phenotype within our vascularized heart-on-a-chip. This necessitated the modification of our platform design to mitigate potential hazard and ensure compatibility with containment level-3 (CL-3) facility working conditions. Briefly, upon SARS-CoV-2 application, PBMCs infiltrated the cardiac tissue from the vascular compartment, creating a hyperinflammatory microenvironment that resulted in electromechanical dysfunction of the cardiac tissue. Impairment of myocardial mitochondria, along with the release of mitochondrial DNA (mtDNA) was found to be a crucial pathophysiological hallmark of the developing myocarditis phenotype. Notably, plasma samples from intensive care unit COVID-19 patients exhibit elevated levels of circulating cell-free mtDNA, correlating with systemic inflammation and inversely associated with cardiac function. HUVEC-EVs imparted a cardiovascular-protective effect and carried immunomodulatory miRNAs that alleviated SARS-CoV-2-induced myocardial injury through toll-like receptor (TLR)-NF-kB axis, which subsequently mitigated the proinflammatory response and mitochondrial impairment. These findings suggest that HUVEC-EVs may serve as a promising immunomodulatory agent for the transfer of anti-inflammatory miRNAs to maintain heart homeostasis and prevent inflammatory injury.

## Results

### Establishing a SARS-CoV-2 induced myocardial inflammation in heart-on-a-chip platform

The InVADE system (**Figure 1**) was leveraged to capture aspects of immune cell activated cardiac tissue dysfunction in the presence of human derived PBMCs. HUVECs were grown inside a hollow biocompatible polymeric vessel (100µm wide x 100µm height) and then cultured with a human iPSC-derived cardiac construct (**Figure 1A**) to generate perfusable functional vascularized heart tissues. Immune cells, PBMCs were perfused through the endothelialized internal lumen of the microfluidic scaffold via a gravity driven flow to investigate how SARS-CoV-2 presence and immune cells interact to aggravate heart function, as well as to screen for the therapeutic benefit of extracellular vesicles (**Figure 1B**). The fluid shear stress exerted within the bioscaffold was estimated to be approximately 1.3 dyne cm^-2^, which closely aligns with the physiological flow conditions experienced by endothelial cells within post-capillary venules^13^, where immune cell infiltration is mostly observed in COVID-19 patients^14–18^. To validate the utility of the InVADE model, SARS-CoV-2 was first introduced into a vascular channel to mimic *in vivo* infection at multiplicity of infection (MOI) 0.1, which is considered physiologically relevant^19^. Following 1.5 h of viral infection, fluorescently labelled PBMCs were perfused into each sample (**Figure 1C**) and samples were subjected to hydrostatic pressure-driven flow. After 72 h of immune cell introduction, DiI stained PBMCs (**Figure 2A and Supplemental Figure 1A**), including CD3+ T-lymphocytes (**Supplemental Figure 1B**), were observed to extravasate from the vascular compartment into the cardiac tissue through the 15µm x 15 µm micro-holes of the microfluidic scaffold. The local accumulation of immune cells was mirrored by increased production of cytokines and chemokines, including IL-6, IL8, and MCP-1 (**Figure 2B and Supplemental Figure 2A**), which further help to attract immune cells toward the site of inflammation guided by chemotactic signals.

**Figure 1:**
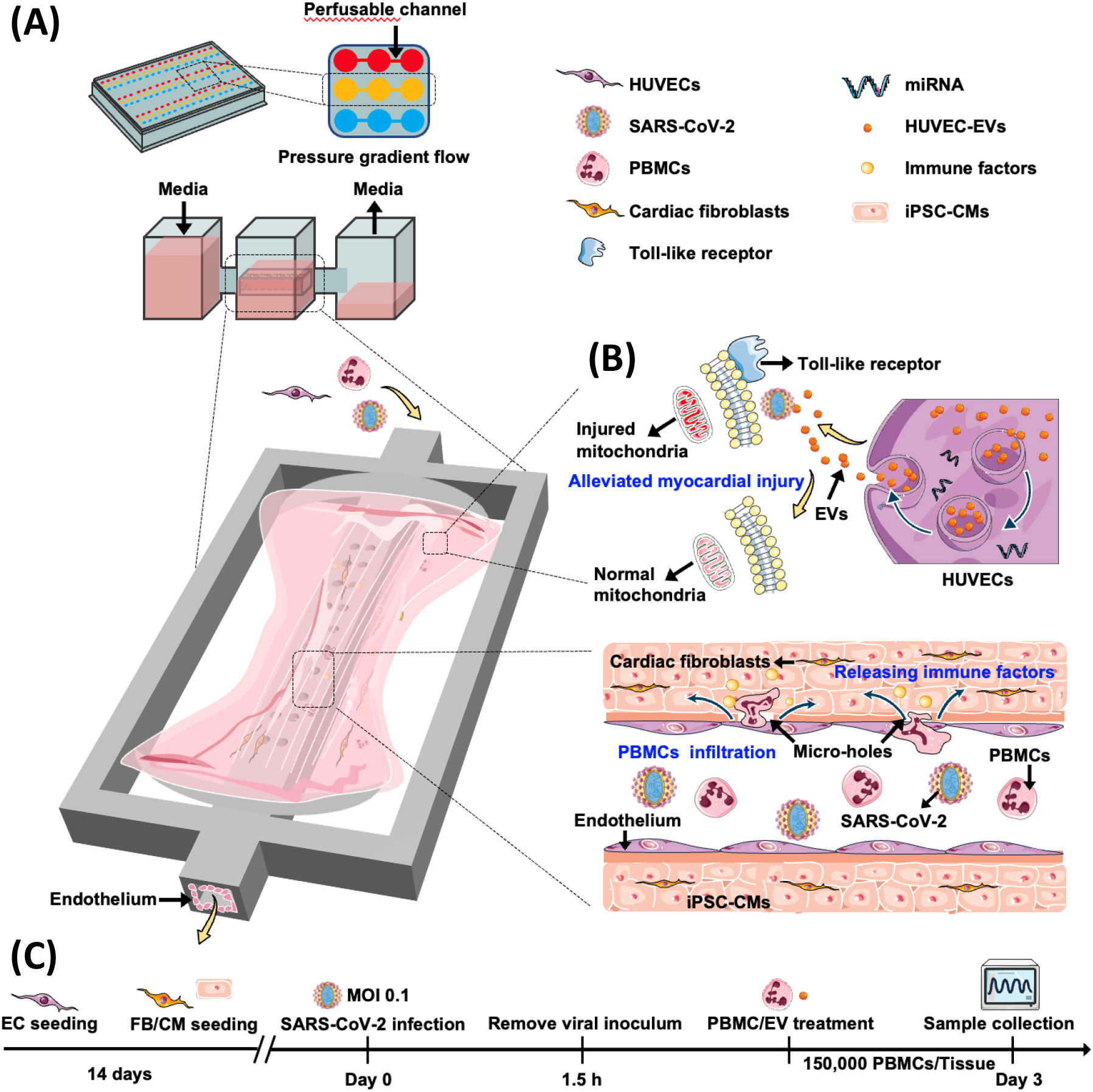
Vascularized heart-on-a-chip model for SARS-CoV-2-induced myocardial inflammation model. (A) Illustration of the components of the InVADE system. Endothelial cells were grown within the luminal space, and iPSC-derived cardiac tissues were cultured within the parenchymal space. Perfusion is initiated through hydrostatic pressure gradient within the vascular compartment. (B) Schematic of the perfusion of human peripheral blood mononuclear cells (PBMCs) through the vasculature. PBMCs can infiltrate into the cardiac tissue compartment through 15µm micro-holes. Extracellular vesicles (EVs) were added to mitigate mitochondrial dysfunction. (C) Experimental timeline showing the sequence of SARS-CoV-2 infection and PBMC addition.

**Figure 2:**
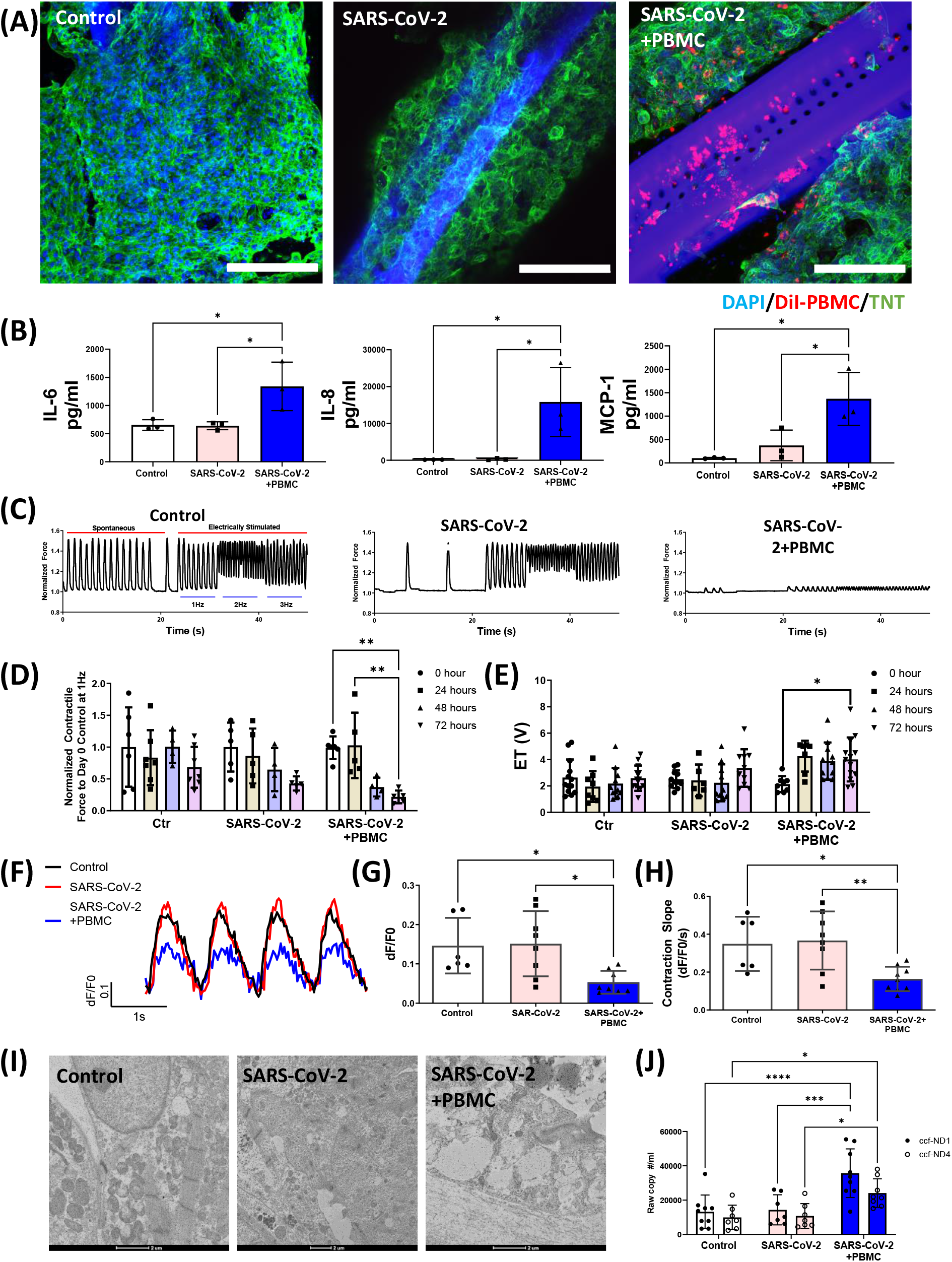
Presence of immune cells in SARS-CoV-2 infected heart-on-a-chip leads to progressive contractile dysfunction in cardiac tissues and the release of cell-free mitochondrial DNA. (A) Representative confocal fluorescence microscopy images of cardiac troponin T disruption in the presence of SARS-CoV-2 and PBMCs. DiI-labelled PBMC (red), nuclei-labelled DAPI (blue), and troponin T (green). Scale bar=200μm. (B) Analysis of proinflammatory cytokines and chemokines 72 h after SARS- CoV-2 infection at MOI 0.1. Data are mean±SD, N=3. One-way analysis of variance (ANOVA) with Bartlett’s test, *p<0.05. (C) Representative trace of cantilever displacement of the cardiac tissue beating paced with increasing stimulation frequency 72 h after SARS-CoV-2 infection at MOI 0.1. (D) Excitation threshold and (E) normalized contraction force amplitude were measured under electrical stimulation at 0, 24, 48, and 72 h after SARS-CoV-2 infection at MOI 0.1. N=3. Two-way ANOVA with Bartlett’s test, *p<0.05, **p<0.01. (F) Representative intracellular calcium transient of cardiac tissue 72 h after infection with SARS-CoV-2 stimulated at 1Hz. Quantification of the intracellular transient properties after 72 h of exposure to SARS-CoV-2: (G) calcium amplitude and (H) contraction slope. Data are mean±SD, N=3. One- way ANOVA with Bartlett’s test, *p<0.05, **p<0.01. (I) Representative TEM images of cardiac tissue sections after infection with SARS-COV-2. Scale bar=2μm. (J) Circulating cell-free mitochondrial transcripts (ccf-mtND1 and ccf-mtND4) in cell culture medium. Data are mean±SD, N=3. One-way ANOVA with Bartlett’s test, *p<0.05, **p<0.01.

### Presence of immune cells leads to a progressive cardiac tissue dysfunction upon SARS-CoV-2 infection

Cantilevers incorporated into the InVADE platform allow for non-invasive monitoring of contractile force using cantilever displacement (**Supplemental Figure 3A**), via previously generated calibration curves (**Supplemental Figure 3B**). Although SARS-CoV-2 presence alone did not induce a contractile deficit in the cardiac tissue, the introduction of PBMCs after SARS-CoV-2 inoculation led to a significant contractile deficit at 72-hour post infection (**Figure 2C** and **Figure 2D**). This was in accord with the findings that highlight the role of infiltrating PBMCs in generating in inflammatory microenvironment in the SARS-CoV-2 infected cardiac tissues, that ultimately culminated in disruption of the contractile proteins, such as troponin T, within the cardiac tissue (**Figure 2A**). The electrical excitability of the cardiac tissue treated with PBMCs progressively declined with increasing viral infection durations, as evidenced by the rise in the excitation threshold (ET), which denotes the minimal voltage required to generate synchronous contraction (**Figure 2E**). Moreover, SARS-CoV-2 inoculation with PBMCs seemingly disrupted intracellular calcium homeostasis (**Figure 2F**), as suggested by the significant reduction in the intracellular calcium transient amplitude (**Figure 2G**) and calcium influx kinetics (**Figure 2H**) as compared to the control group, while the intracellular calcium efflux and decay time constant remained unchanged (**Supplemental Figure 2B**). Importantly, we highlight that the integration of PBMCs in the absence of SARS-CoV-2 does not impinge upon the function of cardiac tissues (**Supplemental Figure 3C and D**). This denotes the cooperative interactions of SARS-CoV-2 with immune cells potentially exacerbates electrochemical regulation of the cardiac tissues. These findings are consistent with our previous study which examined in great detail the introduction of PMBCs alone without SARS-CoV-2 into the endothelialized InVADE platform, documenting no elevation in cytokine secretion or significant immune cell extravasation^20^.

Given that cardiac mitochondria are critical in upholding cardiac homeostasis through their role in providing energy needed for cardiac excitation-contraction coupling and regulating essential intracellular pathways related to cellular survival, the effect of increased inflammation on mitochondrial function was assessed. TEM imaging identified significant mitochondria impairment demonstrated by mitochondrial loss and increased vacuolization in SARS-CoV- 2/PBMC-exposed cardiac tissues (**Figure 2I**). This was also accompanied by elevated levels of ccf-mtDNA. Specifically, ccf-NDA and ccf-ND4, were observed in the culture medium, indicating release of the free-mitochondrial DNA from the cardiac tissue (**Figure 2J**).

### Ccf-mtDNA is a key predictor of cardiac dysfunction in COVID-19 patients

To determine whether an increase in plasma ccf-mtDNA concentration is reflected in COVID-19 patients with cardiovascular complications, assessments were performed in 40 patients admitted to the medical-surgical intensive care units (MSICU) at St. Michael’s Hospital with acute respiratory failure and suspicion of COVID-19 in Toronto, Canada between March 2020 and March 2022. Median patient age was 61.6 years (IQR: 51.3-71.0), ranging from 31 years to 90 years (**Table 1**). Half of the admitted patients were confirmed to be SARS-CoV-2- positive by polymerase chain reaction tests. On average, patients with COVID-19 (n=20) had higher ccf-mtND1 (median 11102 copies/ml [IQR: 5988-27885]) and ccf-mtND4 (median 8849 copies/ml [IQR: 5408-20614]) levels compared to SARS-CoV-2-negative patients (n=20) (ccf- mtND1 median 4463 copies/ml [IQR: 1392-12385], p=0.0316; ccf-mtND4 median 3694 copies/ml [IQR: 1049-7692], p=0.00246) (**Table 1**). Of note, higher ccf-mtDNA levels were measured in COVID-19 patients with left ventricular ejection fraction (LVEF) <50% compared to COVID-19 patients with LVEF≥50% (**Figure 3A**). Higher mortality rates were documented for the former compared to the latter group of patients (70% vs. 20%; **Figure 3A**). Within the COVID-19 patient cohort, a significant negative association was found between plasma ccf-mtDNA levels with LVEF (**Figure 3B**), suggesting that elevated concentration of ccf-mtDNA is closely linked to increased risk of cardiac dysfunction. Furthermore, a direct linear correlation between ccf-mtDNA levels and systemic inflammation was observed (**Figure 3B**), consistent with clinical observations in patients experiencing COVID-19 cardiac injury^21^. Taken together, these results establish that heart-on-a-chip system with circulating immune cells is capable of pinpointing potential biomarkers which closely align with clinical observations. Notably, the plasma concentration of ccf-mtDNA emerges as a robust marker indicative of both cardiac dysfunction and systemic inflammation and may facilitate early identification of patients with poor prognosis.

**Figure 3:**
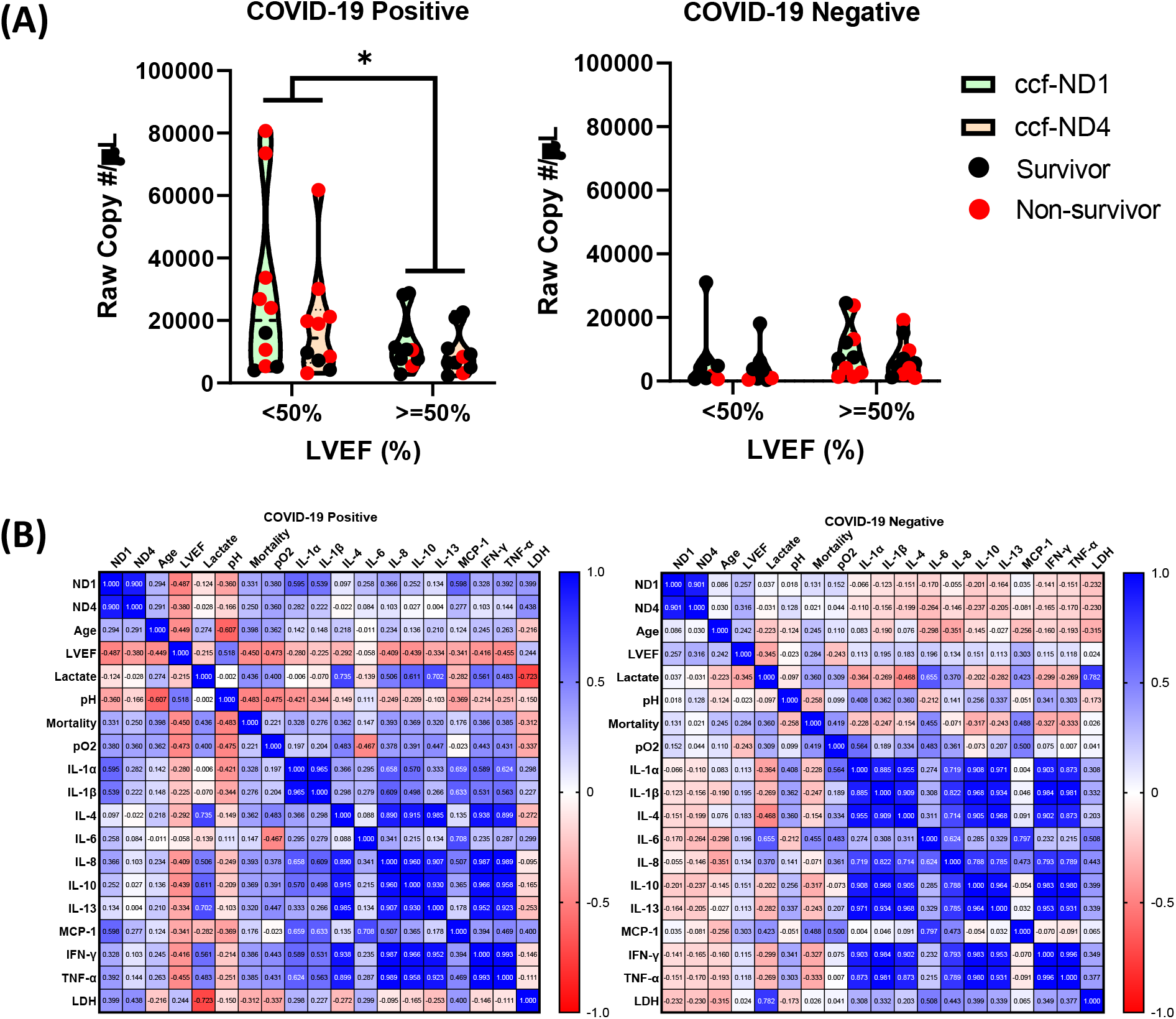
Circulating cell free mitochondrial DNA (ccf-mtDNA) is elevated in severe COVID-19 patients with contractile dysfunction. (A) Concentration of ccf-mtDNA in patient plasma samples. Data are mean±SD, n=20. Two-way ANOVA with Bartlett’s test, p<0.05. (B) Pearson correlation of patient characteristics and plasma concentration of ccf-mtDNA within the COVID-19-positive and COVID-19- negative subgroups.

**Table 1:** Summary of the clinical characteristics and age of ICU-admitted patients.

| Clinical Characteristics |  |  |  |
| --- | --- | --- | --- |
|  | COVID-19 + (n=20) | COVID-19 - (n=19) |  |
| Demographics |  |  | p-value |
| Age, median (IQR)- yr. | 62.5 (49.25-72.25) | 67.0 (51.25-70.0) |  |
| Distribution no. (%) |  |  |  |
| 18-40 yr. | 2 (10) | 2 (10) |  |
| 41-64 yr. | 9 (45.0) | 6 (30) |  |
| >=65 yr. | 9 (45.0) | 12 (60) |  |
| Clinical Findings |  |  |  |
| Lactate (IQR) – mmol/L | 1.40 (1.10-1.1.80), 13/20 | 1.70 (1.20-2.30), 11/20 | 0.3146 |
| pO2 (IQR) - mmHg | 77 (49.75-88.75), 16/20 | 85 (65.0-120.5), 17/20 | 0.0868 |
| pH (IQR) | 7.375 (7.310-7.435), 16/20 | 7.37 (7.315-7.415), 17/20 | 0.9187 |
| Laboratory Findings |  |  |  |
| Total ccf-mtDNA ND1 (IQR)<br>– count/μl<br>All patients | 11102 (5988-27885) | 4463 (1392-12385) | 0.0316 |
| Patients with LVEF<50 | 17319 (4920-43644) | 2879 (742-6329) | 0.0782 |
| Patients with LVEF≥50 | 10650 (7107-19635) | 7238 (2369-15832) | 0.7328 |
| Total ccf-mtDNA ND4 (IQR)<br>– count/μl<br>All patients | 8849 (5408-20614) | 3964 (1049-7694) | 0.00246 |
| Patients with LVEF<50 | 13737 (3968-23424) | 2136 (559-5630) | 0.0656 |
| Patients with LVEF≥50 | 6516 (5046-11052) | 4565 (1696-14398) | 0.7255 |

### HUVEC-derived EVs as a potential therapy for SARS-CoV-2-induced contractile dysfunction and ccf-mtDNA release

As inflammatory cascade starts off with endothelial cell and immune cell activation, we hypothesized that EVs derived from quiescent endothelial cells may be crucial in balancing the inflammatory response, given their significant contribution to the preservation of vascular homeostasis. EVs were isolated from quiescent HUVECs using precipitation method, and they displayed a cup-shaped morphology with an average diameter of 150 nm (**Figure 4A**). Nanoparticle tracking analysis revealed that EVs had a particle size of 155nm, which was comparable in size to the TEM image (**Figure 4B**). Western blot analysis confirmed the presence of exosomal markers, including CD9, CD63, and flotillin-1. In parallel, no intracellular protein calnexin was detected, indicating pure EV isolation (**Figure 4C**).

**Figure 4:**
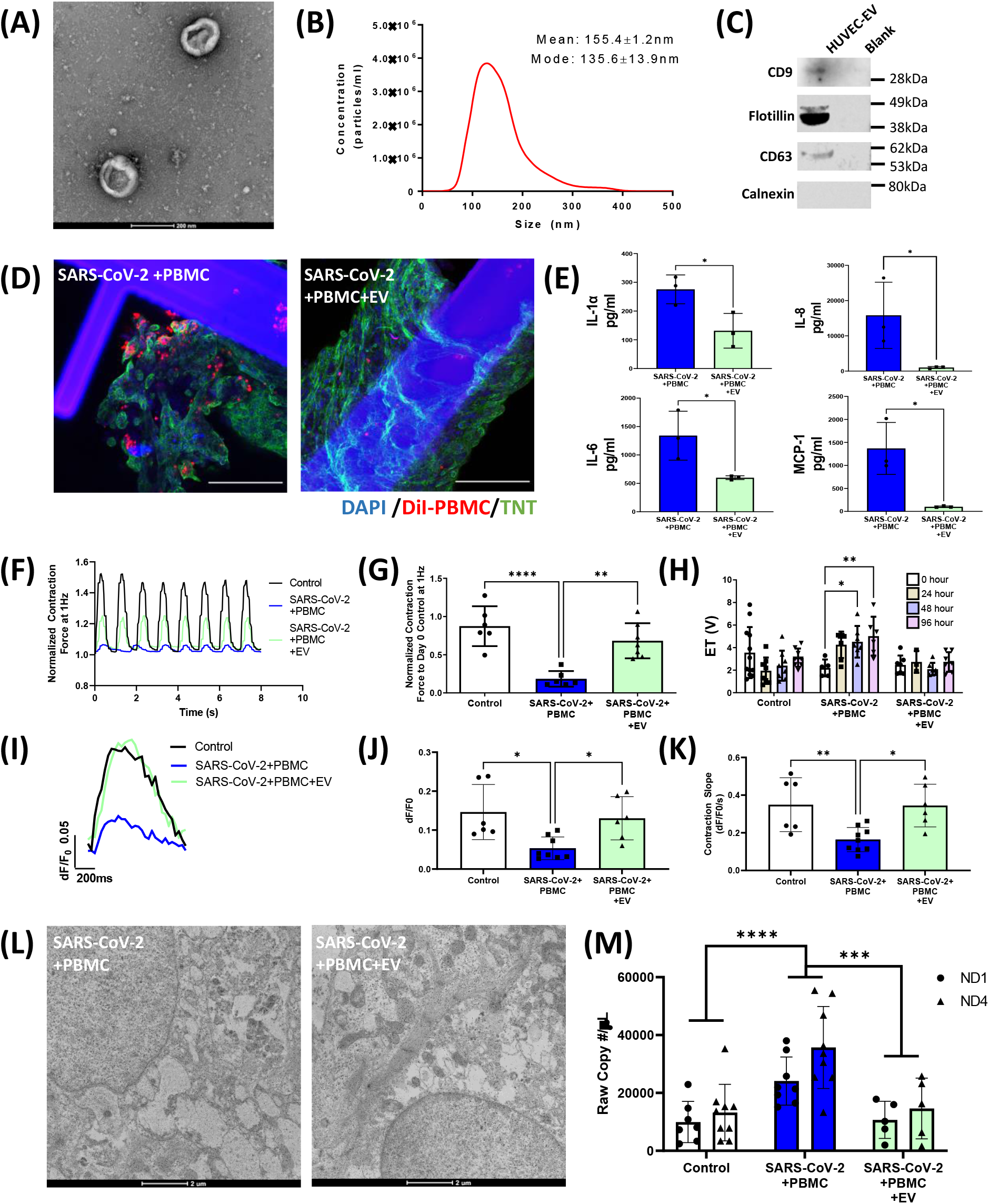
Extracellular vesicles promote recovery of cardiac tissue functions following SARS-CoV-2 application in the presence of immune cells and mitigate the release of cell-free mitochondrial DNA. (A) Representative TEM image of HVUEC-EVs. Scale bar = 200nm. (B) Representative size distribution of HUVEC-EVs as measured by nanoparticle tracking analysis. (C) Immunoblots of HUVEC-EVs for the exosomal markers CD9, CD63, and Flot-1. The absence of calnexin demonstrates pure EV isolation (D) Representative confocal fluorescence microscopy images of PBMCs infiltrating the human iPSC-derived cardiac tissue upon HUVEC-EV treatment. DiI labelled PBMCs (red) and nuclei-labeled DAPI (blue). Scale bar=100μm. (E) Analysis of proinflammatory cytokines and chemokines from cardiac tissue 72 h after SARS-CoV-2 infection at MOI 0.1. Data are mean±SD, N=3. Student’s t-test with Bartlett’s test, *p<0.05, **p<0.01. (F) contractility. (G) Normalized contraction amplitude and (H) excitation threshold were measured under electrical stimulation at 1Hz, 72 h after SARS-CoV-2 infection at MOI of 0.1. (I) Representative TEM images of cardiac tissue after SARS-CoV-2 infection. Scale bar = 2μm. (J) Concentration of ccf-mtDNA in cell culture medium collected 72 h after SARS-CoV-2 exposure and treatment with HUVEC-EVs. Data are mean±SD, N=3 from each experiment. Two-way ANOVA with Bartlett’s test, *p<0.05, **p<0.01, 11 ***p<0.001, ****p<0.0001.

When co-administered with PBMCs in the vascular compartment, EVs led to a decrease in the infiltration of PBMCs into the cardiac tissue (**Figure 4D**). This was accompanied by a reduction in the measured proinflammatory cytokine responses, namely IL-1α, IL-6, IL-8, and MCP-1 (**Figure 4E**), and a notable shift towards anti-inflammatory phenotype, characterized by increased IL-10 levels (**Supplemental Figure 4A**). A significant improvement in the contractile function of the SARS-CoV-2-infected heart-on-a-chip were observed (**Figure 4F and G**), leading to the restoration of electrical excitability properties to the levels comparable to those of control tissues (**Figure 4H**). Furthermore, the amplitude (**Figure 4I and J**) and kinetics of intracellular calcium transient (**Figure 4K; Supplemental Figure 4B-E**) were restored to normal levels, as were the sarcomere structure and mitochondrial ultrastructure (**Figure 4L**). In addition, a decrease in ccf-mtDNA release into the extracellular environment were measured. Collectively, these results support the notion that HUVEC-EVs can serve as a promising therapy for restoration of mitochondrial integrity and repairing myocardial damage in the context of SARS-CoV-2 inflammation.

### Extracellular vesicles contain miRNAs that suppress cellular proinflammatory responses

A microRNA sequencing (miRNA-seq) analysis of EV cargo was performed in search of miRNAs that regulate gene expression (**Supplemental Data 1**). A total of 561 unique miRNAs were detected in isolated EVs (**Figure 5A**) and ranked according to fraction of total mapped miRNA reads. The top-9 abundant miRNA in EVs were let-7b-5p, miR-1246, miR-126-3p, let- 7a-5p, let-7c-5p, let-7i-5p, let-7f-5p, miR-16-5p, miR-125b-5p, and let-7e-5p (**Figure 5B**). Subsequently, a gene ontology (GO) enrichment analysis was run to identify the biological processes that were significantly upregulated in the dataset. The targeted pathways of the top-9 most abundant miRNAs primarily clustered to biological processes relating to the toll-like receptor (TLR) singling pathway, immune response-regulating signaling pathway and mitogen-activated protein kinase (MAPK) cascade (**Figure 5C**). A Kyoto Encyclopedia of Genes and Genomes (KEGG) analysis found EV target signaling pathways associated with the TLR and tumor necrosis factor (TNF) signaling pathways (**Figure 5D and E**). As miRNAs typically repress mRNA function, this finding suggests that HUVEC-EVs contain miRNAs that help improve the heart function by repressing TLR activation and subsequent inflammation induced by SARS-CoV-2 infection in endothelial cells, immune cells and cardiomyocytes.

**Figure 5:**
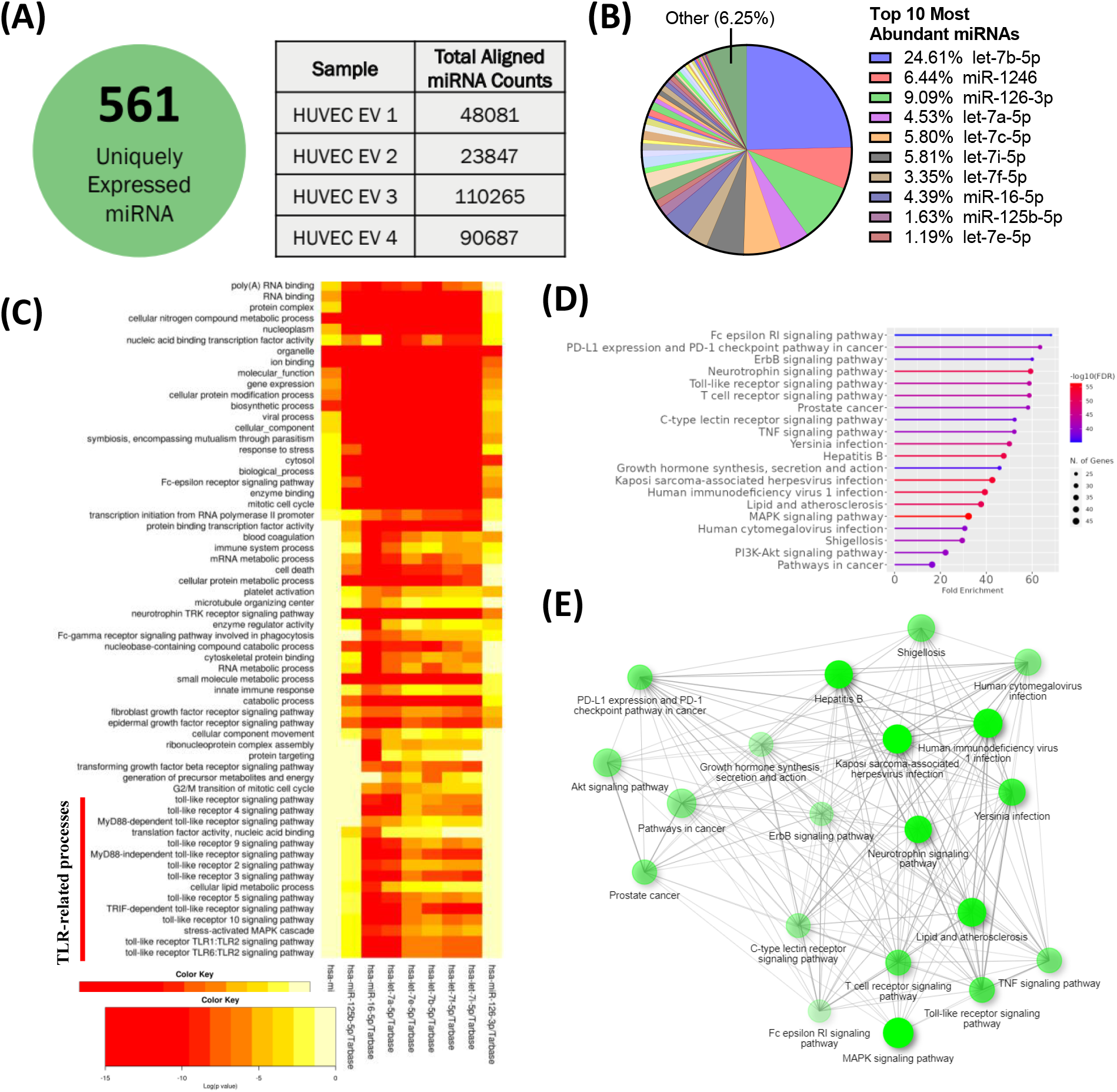
miRNA sequencing reveals that extracellular vesicles contain microRNAs that target TNF and TLR signaling pathways. (A) A diagram demonstrating the amount of uniquely expressed miRNAs in the HUVEC-EVs and their total aligned miRNA counts. (B) Fraction of 50 most-abundant miRNAs in HUVEC-EVs. (C) Heatmap of enriched biological processes for top-9 most-abundant miRNA-derived from HUEV-EVs. (E) KEGG pathway enrichment analysis for top-9 most-abundant miRNAs in HUVEC-EVs. Y-axis indicates the pathway name and x-axis indicates the enriched factors in each pathway. Bubble size indicates the number of genes. (F) the interaction of top-20 significantly enriched KEGG pathways for top- 9 abundant miRNAs in HUVEC-EV.

To validate these findings, TNF signaling was assessed following EV treatment of TNF- α-stimulated endothelial cells (**Figure 6A**). After 2 h of transfection, EVs were detected within HUVECs (**Figure 6B**) along with an approximately 6-fold increase in *TNF*α, a 3-fold increase in *IL8* mRNA and suppression of proinflammatory genes (**Figure 6C**). These results provide compelling evidence that EVs can mitigate inflammatory responses, as predicted by the GO and KEGG analyses.

**Figure 6:**
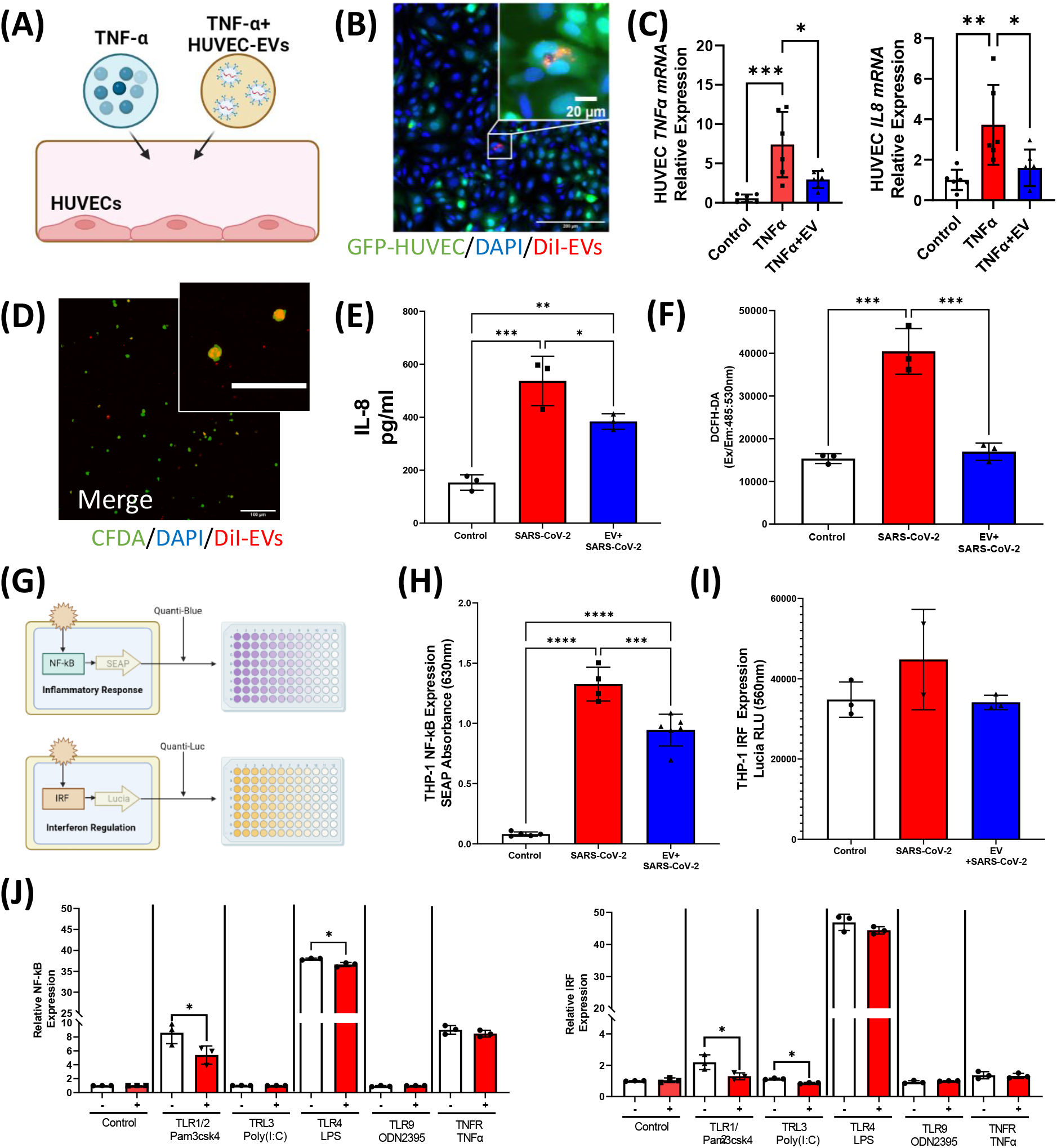
Extracellular vesicles provide anti-inflammatory and cardioprotective effects through the toll-like receptor (TLR)-NF-kB axis. (A) Schematic of experiments. HUVECs were stimulated with either TNF-α or TNF-α/HUVEC-EVs. (B) Representative confocal microscopy image of DiI-3 stained (red) EVs inside GFP-HUVECs. Scale bar = 200μm. Magnified image scale bar = 20μm. (C) Quantification of *TNFα* and *IL8* gene expression in HUVEC. Data are mean±SD, n=6. One-way ANOVA with Bartlett’s test, *p<0.05, **p<0.01, ***p<0.001. (D) Representative confocal images of DiI-stained HUVEC-EVs (red) inside PBMCs (green: CFDA) Scale bar =100μm. Magnified image scale bar =50μm. (E) Analysis of proinflammatory IL-8 secreted by PBMCs 24 h after SARS-CoV-2 infection at MOI 1. Data are mean±SD, n=3. One-way ANOVA with Bartlett’s test, *p<0.05, **p<0.01, ***p<0.001. (F) Extracellular ROS level was measured using the ROS-sensitive colorimetric dye, 2’7,-dichlorofluorescein diacetate (DCFH-DA) at 530nm. Data are mean±SD, N=3. Student’s t-test with Bartlett’s test, *p<0.05, **p<0.01, ***p<0.001. (G) Illustration depicting dual reporter THP-1 cells used to monitor the NF-kB pathway, by monitoring the activity of SEAP, and the IRF pathway, by assessing the activity of a secreted luciferase, Lucia. THP-1 cells were treated for 24 h with SARS-CoV-2 at MOI 1. Cell culture medium were assessed for (H) SEAP reporter activity and (I) Lucia reporter activity. Data are mean±SD, n=3. One-way ANOVA with Bartlett’s test, *p<0.05, **p<0.01, ***p<0.001. (J) Cells were stimulated with 1ng/ml Pam3CSK (TLR1/2), 5μM poly(I:C) (TLR3), 10ng/ml LPS (TLR4), 5μM ODN2395 (TLR9) or 1ng/ml TNF-α (TNFR). After a 24 h incubation, NF-kB and IRF activation were assessed by measuring the levels of SEAP and Lucia, respectively. Data are mean±SD, N=3. Student’s t-test with Bartlett’s test, *p<0.05.

### Extracellular vesicles target the TLR-NF-kB axis to mitigate proinflammatory responses

To characterize the effect of HUVEC-EVs on SARS-CoV-2-mediated proinflammatory response, PBMCs were treated with EVs and then stimulated with SARS-CoV-2. Uptake of EVs by PBMCs, confirmed by DiL-labeling of the EVs (**Figure 6D**), resulted in the suppression of IL-8 activation (**Figure 6E and Supplemental Figure 5**), providing further evidence that HUVEC-derived exosomes can blunt recruitment of immune cells to the site of inflammation (**Figure 6E**). Additionally, levels of extracellular reactive oxygen species (ROS) were significantly lowered following EV treatment, suggesting that EVs can protect mitochondrial components from damage and preserve their function (**Figure 6F**).

We then used THP-1 dual-reporter cells, a validated monocyte cell model, to examine whether EVs can regulate NF-kB, a central regulator of innate immunity. THP-1 dual-reporter cells secrete inducible reporter proteins Lucia and secreted embryonic alkaline phosphate (SEAP), which reflect the activation of interferon-related factors (IRF) and NF-kB, respectively (**Figure 6G**). THP-1 cells stimulated with SARS-CoV-2 secreted significantly more SEAP than non-stimulated cells, confirmed that SARS-CoV-2 infection activate NF-kB signaling (**Figure 6H**). In contrast, IRF expression was not changed upon SARS-CoV-2 infection, consistent with the finding that SARS-CoV-2 antagonizes the interferon responses IFNAR1 lysosomal degradation (**Figure 6I**) ^22, 23^. Treatment with EVs prior to SARS-CoV-2 infection limited NF-kB activation (**Figure 6H**), demonstrating that HUVEC-EVs mitigate inflammatory responses by inhibiting the NF-kB cascade.

A thorough screening of various innate sensors located upstream of NF-kB was then conducted to identify the specific EV-target receptors. NF-kB expression was significantly increased in THP-1 cells challenged with Pam3csk4 (agonist of TLR1 and TLR2 heterodimer), lipopolysaccharide (LPS; TLR4 agonist) or TNF-α (agonist of TLR) (**Figure 6J**). Conversely, THP-1 cells primed with EVs prior to the addition of TLR agonist or TNF-α, showed reduced Pam3csk and LPS-induced NF-kB activation, but similar levels of TNF-α-mediated NF-kB activation (**Figure 6J**). Moreover, EVs were unable to supress NF-kB activation when THP-1 cells were stimulated with individual cytokines (TNF-α, IL-6, IFN-β) or a cocktail of cytokines (**Supplemental Figure 6**), suggesting that EVs primarily target TLR-initiated inflammation and downstream factors. Collectively, these data show that EVs regulate proinflammatory responses of monocytes and shift activation toward inflammatory resistance state, thereby mitigating local and systemic damage.

### Extracellular vesicles reverse inflammation-induced cardiac dysfunction

Clinical data have demonstrated that patients infected with SARS-CoV-2 have elevated levels of IFN-γ, IL-6, and TNF-α (**Supplemental Figure 7**) ^24, 25^. This cytokine cocktail was applied here to a cardiac bundle termed Biowire in the absence of endothelial and immune cells, to benchmark direct effects of EVs on restoration of contractile function (**Supplemental Figure 8A**). The application of cytokine cocktail led to a modest reduction in contractile force (**Supplemental Figure 8B**) and maximum capture rate (**Supplemental Figure 8C**), and to an increase in excitation threshold (**Supplemental Figure 8D**), all of which correlate with cardiac dysfunction. Importantly, HUVEC-EVs rescued dysfunctional cardiac tissues, as illustrated by a recovery of contraction force (**Supplemental Figure 8B**) and tissue excitability (**Supplemental Figure 8D**). Collectively, these data show that EVs can both reduce the magnitude of proinflammatory responses and contribute to directly preserve cardiomyocyte function in the inflammatory environment.

## Discussion

High fidelity, multi-cell type, human *in vitro* models of inflammatory conditions of the myocardium, remain scarce, ultimately slowing the progress towards new therapeutics. Considering COVID19 as an example, among the 43 studies reviewed by Haussner *et al.*, treatment for COVID-19-associated myocarditis was highly variable, with low utilization^26^. Cardiovascular involvement was first recognized in China, where patients with elevated serum levels of troponin I, IL-6 and other pro-natriuretic peptides were found to have an increased risk of mortality^24^. Most therapeutic options still rely on the use of ACE inhibitors or beta blockers, while treatments focusing on the immune system and subsequent systemic inflammatory response are scarce. Here, we developed a model of human SARS-CoV-2 induced myocarditis on-a-chip, demonstrating disease hallmarks such as the increased levels of cytokines triggered by the viral presence, which, in turn, further amplify cytokine release and the recruitment of circulating immune cells, thereby accelerating progression of cardiac dysfunction ^27–29^.

Previously, we demonstrated that endothelial cells switch to a proinflammatory phenotype upon SARS-CoV-2 infection in the InVADE platform, as manifested by elevated release of soluble vascular adhesion molecules (E-selectin and ICAM-1) and proinflammatory cytokines (angiopoietin-2, IL-6, IL-8, and TREM-1)^20^. This endothelial cell activation promotes monocyte recruitment, leading to the activation of immune cells. The present study demonstrated that inflammation can also lead to the infiltration of monocytes and CD3 T-lymphocytes, via the endothelium, into the cardiac parenchyma. This is consistent with the increased CD3+ T-lymphocyte levels measured post-mortem in the heart tissues of COVID-19 patients with myocarditis^14–17^. Local infiltration of the immune cells results in an acute inflammatory response, causing the release of IL-8 and MCP-1 which further promotes monocyte recruitment to the site of inflammation. This, in turn impairs cardiac function, consistent with the findings that immune cells are the primary cause of cardiac injury in COVID-19 patients^14, 17, 30^

The heart is one of the most metabolically demanding organs and requires a large number of mitochondria to produce ATP. Here, excessive myocardial inflammation increased ccf-mtDNA presence, namely ccf-mtND1 and ccf-mtND4 in the heart-on-a-chip model, which was also observed in patients diagnosed with COVID-19 admitted to the ICU with low LVEF (<50%). Several clinical studies found that patients with higher ccf-mtDNA concentrations are more likely to require intensive care and higher mortality^31, 32^. Therefore, we suggest that ccf-mtDNA could be used as a biomarker to identify COVID-19+ patients at risk of more serious LVEF dysfunction and predict clinical outcomes in COVID-19+ ICU patients. Our findings are consistent with previous studies, which report that in the setting of cardiac injury, elevated levels of intracellular ROS triggered by proinflammatory cytokine activation led to mitochondrial membrane permeabilization and subsequent cell death^33^. This cascade of events culminates in reduced expression of mitochondrial genes involved in oxidative phosphorylation, ATP metabolism and mitochondrial transition^19^, ultimately leading to mitochondrial dysfunction. The impaired mitochondria then release mtDNA as well as mitochondrial proteins^34^, which may act as danger-associated molecular pattern DAMPs that activates immune responses and elicit further damage^35, 36^. Various inflammatory pathways, such as cGAS-STING and TLR4/9 and inflammasome formation, can lead to robust type I IFN responses and NF-kB-dependent proinflammatory responses, which may set off the apoptotic cascade^37^.

These redundant and reinforcing inflammatory interactions in viral myocarditis pose a significant challenge in pinpointing the critical cytokines/chemokines that drives the phenotype, necessitating a comprehensive strategy for inhibiting cytokine release. Several anti-inflammatory agents have shown clinical benefits in the acute management of COVID-19. For example, bromodomain and extraterminal family inhibitors (BETi) improved cardiac function in the lipopolysaccharide-induced mouse cytokine-storm model; however, loss of BET activity may negatively alter energy production in the mitochondria^38^. Corticosteroids, such as dexamethasone, may also improve patient mortality, but systemic immunosuppression can impede viral clearance, thereby potentially exacerbating the disease^39–42^.

Taken together, there remains a need for immunomodulatory compounds that can attenuate cytokine-induced cardiac dysfunction while preserving the mitochondrial energetics in patients with SARS-CoV-2-induced myocarditis. As endothelial cell-derived EVs have been shown to supress immune cell activation^12^ and alleviate cardiac cell death^10^, this study tested the immunomodulatory potential of HUVEC-derived EVs to alleviate SARS-CoV-2-induced myocardial inflammation. HUVEC-EVs improved cardiac functionality after SARS-CoV-2 infection, as demonstrated by the recovery of electro-mechanical properties and mitigated release of ccf-mtDNA. miRNA-sequencing identified let-7 family, miR-16 and miR-125b as the most abundant miRNA in HUVEC-EV cargo, which were predicted to target apoptotic processes, TLR signaling and immune system processes. Additionally, KEGG analysis demonstrated that the EVs target signaling pathways associated with TLR and TNF signaling pathway. Thus, EVs may provide therapeutic benefits by targeting genes associated with inflammation, thereby mitigating the production of proinflammatory cytokines. Indeed, EV treatment of THP-1 reporter cells with monocyte-like phenotype mitigated NF-kB, a central downstream transcriptional factor that governs inflammation and cellular stress, supressed the TLR-mediated NF-kB activation observed following SARS-CoV-2 infection. Importantly, this inhibition reduced the release of proinflammatory cytokines (IL-8) as well as mRNA level of *Il-8* and *Tnf-*α, thereby impeding the activation of NF-kB, and prevented positive feedback between cytokines and circulating monocytes. Given that TLR signaling is one of the enriched targets of HUVEC- EVs, the inhibition of this step is the preferred route to protect the heart against inflammatory injury.

Our results substantiate the hypothesis that circulating immune cells are probable contributors to myocardial dysfunction in the context of COVID-19. Nevertheless, the role of resident macrophages in contributing to cardiac pathology associated with this disease remains a significant consideration and requires more understanding. Additional limitations of this work include a relatively short observation time points, absence of microvasculature in the parenchymal space, the need to a more precise and controlled inclusion of adaptive immunity and a systematic testing of the effects of the identified miRNAs.

## Conclusion

COVID-19 is associated with an array of disease manifestations, including a massive inflammatory response that causes myocarditis. Here, we developed a vascularized heart-on-a-chip model of SARS-CoV-2 induced myocarditis to support therapeutics discovery and mechanistic studies. This work showed a tight correlation between elevated ccf-mtDNA and cardiovascular dysfunction in both a human engineered myocarditis model and patients with COVID-19. The increase was traced back to proinflammatory mediators stimulated by TLR pathway. Therefore, a novel therapeutic option targeting TLR may present an important and viable approach to reduce SARS-CoV-2-induced myocarditis and to improve patient outcomes. The ability of HUVEC-EVs to inhibit TLR-mediated NF-kB activation, thereby reducing myocardial inflammation, mitochondrial damage and improving cardiac function, was demonstrated. Future studies characterizing the activities of miRNAs in HUVEC-EVs will help determine effective means of deactivating SARS-CoV-2-induced inflammation.

## Acknowledgements

We would like to acknowledge the training, support and expertise provided by the management of C-CL3 Unit at the Temetry Faculty of Medicine, University of Toronto. Operational support for the C-CL3 Unit throughout the pandemic was provided by the Temetry Foundation and the Toronto COVID Action Fund (TCAI). A. C. A. was supported by Canada Research Chair which supported the ccf-mtDNA analysis. This work was funded by the National Institutes of Health Grant 2R01 HL076485, Natural Sciences and Engineering Research Council of Canada (NSERC) Discovery Grant (RGPIN 326982-10), NSERC Strategic Grant (STPGP 506689-17), Canadian Institutes of Health Research (CIHR) Foundation Grant FDN-167274, Canada Foundation for Innovation/Ontario Research Fund grant 36442. M.R. was supported by Canada Research Chairs and Killam Fellowship. R. X. Z. L. was supported by Alexander Graham Bell Canada Graduate Scholarship-Doctoral Award and PRIME Fellowship. Y. Z. was supported by CIHR Post-Doctoral Fellowship. The COVID-19 Longitudinal Biomarkers of Lung Injury study (COLOBILI) was funded by the St. Michael’s Foundation, Immune Task Force Grant and CIHR grant (GA4-177735) to CCDS and AJB. CCDS is supported by the CIHR (MOP-130331, MOP- 106545, CIHR/NSERC MOP-510282 2020) and the University of Toronto Robert and Dorothy Pitts Research Chair in Acute Care and Emergency Medicine. We would like to thank Dr. Stephen Juvet for giving access to the equipment for cytokine quantification.I (MR) would like to dedicate this study and this manuscript to my grandmother who passed away in the ICU from COVID19 induced heart failure. Thank you for your love and encouragement.

## Author contribution

R. X. L., N. R., Y. Z., and M. R. designed the study. R. X. L. and Y. W. generated the device and tissue constructs. N. R. performed all the experiments involving SARS-CoV-2, and R. X. Z. performed the subsequent analyses. K. T. W. performed EV-NTA and miRNA sequencing. E. L. B. conducted the ccf-mtDNA and extracellular ROS analysis. C. C. S., U. T., A. B. and E. C. prepared and provided patient plasma samples. J. C. and R. X. Z. L. performed all experiments associated with the THP-1 cells. D. V. and R. X. Z. L. performed cytokine array experiments. R. X. Z. wrote the initial manuscript, which was edited by N. R., K. T. W., A. C. A., B. L., E. L. B., Y. Z., C. C. S. and M. R. All authors read and approved the manuscript.

## Materials and Methods

### InVADE platform fabrication

InVADE platform was prepared as we described previously ^27–29^. Briefly, the fabrication of the silicon master mold involved the use of the standard soft lithography technique, with photomasks designed using AutoCAD software. To prepare the polydimethylsiloxane (PDMS) molds, a mixture of 10 parts silicon elastomer and 1 part curing agent (Sylgard 184, Dow Corning) was prepared. PDMS molds were made by replica molding from the SU-8 mold. Subsequently, a biocompatible UV-crosslinkable elastomeric polymer known as poly(octamethylene maleate (anhydride) citrate) (POMaC), consisting of a combination of 1,8-octanediol, citric acid, and maleic anhydride, mixed with poly(ethylene glycol) dimethyl ether (PEGDM) porogen at a 6:4 (POMaC:PEGDM, wt/wt) ratio, along with 5% 2-hydroxy-1-[4(hydyroxyethoxy)phenyl]-2- methyl-1-propane (Irgacure 2959; Sigma-Aldrich) photoinitation, was perfused through PDMS channels of the mastermold. Through exposure to a UV light source, scaffolds with a square luminal structure and lid were crosslinked. The top and bottom features were then bonded together using a 3D stamping technique as we previously described ^27–29^, to create a perfusable microvessel, termed Angitube patterned with microholes (15µm x 15µm) spaced at 15µm apart as we described^28^. The Angiotube scaffolds with inner luminal dimension of 100µm and wall thickness of 50µm were then placed onto a patterned hot-embossed polystyrene base plate that had a foot-print of a 96-well plate, and the plate was bonded onto a bottomless 96-well plate using polyurethane glue (GS Polymers). By fitting the Angiotube microvessel into groves of the patterned hot-embossed base, it is possible to connect three wells in one column of a 96-well plate. The three wells become a unit for cell cultivation. Before starting cell culture experiments, the entire plate underwent sterilization by treating it with 70% filtered ethanol for 2 h at room temperature.

### Cell Culture

HUVECs were cultured and maintained in endothelial cell growth medium (EGM-2, PromoCell) under standard cell culture conditions, with 5% CO_2_ at 37°C. To endothelialize the polymeric lumen of the InVADE system, 0.2% w.t. bovine gelatin (Type A; Sigma-Aldrich) was coated for 1.5 h onto the POMAC microvessel, followed by overnight EGM-2 medium conditioning. For endothelialization of the Angiotube, 5µL of concentrated endothelial cell suspension (25x10^6^ cells/ml) were seeded at both inlet and outlet, allowing endothelial cells to be packed inside the lumen for 1.5 h at 37°C. Unattached endothelial cells were then flushed by adding EMG-2 media into the inlet to initiate perfusion.

Ventricular cardiomyocytes were derived from human iPSC line BJ1D using a monolayer differentiation protocol, as previously described^43^. To generate cardiac tissue on the InVADE platform or Biowire system described below, human iPSC-derived cardiomyocytes were suspended in fibrinogen from human plasma (Sigma-Aldrich) at a concentration of 50x10^6^ cells/ml. Then, 1µL thrombin from human plasma was added to 3.5µL of the cardiomyocyte- human fibrinogen mixture (Sigma-Aldrich). Then, 3µL of the mixture containing cardiomyocytes was placed onto the InVADE bioscaffold. The cardiomyocyte mixture was then allowed to undergo crosslinking for 10 min at 37°C. The following medium was used for cardiac tissue maintenance: Induction 3 Medium (I3M): stepPro-34 media serum-free medium (Gibco), 20x10^-3^M HEPES (Gibco), 1% GlutaMAX (Gibco), 1% penicillin-streptomycin (Gibco) and 213µg/ml 2-phosphate ascorbic acid (Sigma-Aldrich) containing aprotinin from bovine lung (1:1000 dilution) (Sigma-Aldrich). Perfusion in this system is achieved by hydrostatic pressure gradient, in which 500µL of EGM-2, 350µL of I3M, and 20µL of EGM-2 media were added to inlet, tissue chamber and outlet of InVADE system, respectively. The continuous flow in the scaffold is maintained by placing the InVADE system on a rocker (Perfusion Rocker Mini, MIMETAS) with 20° tilt that is automated to reverse the flow every 4 h. The endothelial media perfuses the microvessel by moving between the inlet and outlet well via a gravity driven flow. This is estimated to achieve a flow rate of 1.4µL/min and a shear stress of 1.3 dyne/cm^2^ according to our previous studies^20, 28^. The tissue well contains cardiac media to support iPSC derived cardiomyocytes.

Human peripheral blood mononuclear cells were obtained from STEMCELL Technologies (cat#700.25.1). PBMCs were cultured and maintained in RPMI1640 with 10% fetal bovine serum (FBS) and 1% penicillin-streptomycin under standard cell culture conditions, with 5% CO_2_ at 37°C.

THP-1 dual-reporter cells (InvivoGen) were cultured and maintained in RPMI1640 with 10% heat-inactivated FBS, 1% normocin and 1% penicillin-streptomycin under standard cell culture conditions, with 5% CO_2_ at 37°C.

### Viral Infection

All handling of SARS-CoV-2/SB2 was conducted in the combined containment level 3 (C-CL3) unit at the University of Toronto. SARS-COV-2 was expanded in Vero E6 cells. Viral dosing was defined by TCID50 in Vero E6 cells. Before the viral infection, both endothelial cell compartment and the cardiac tissue compartment were washed with serum free EMEM media, then media was replaced with serum free EMEM containing SARS-CoV-2 at MOI of 0.1 and perfused through the endothelial cell compartment for 90 minutes. Virus containing media was replaced by EGM2 and I3M media for the endothelial cell compartment and cardiac tissue compartment, respectively. PBMCs were suspended in EGM-2 media at concentration of 300,000 cells/ml, and 500µL of EGM-2 containing PBMCs was applied by perfusion from the inlet for the InVADE experiment after viral infection.

### Immunostaining

SARS-CoV-2-infected samples were fixed with 10% formalin for 1 h at room temperature, as recommended by C-CL3 safety protocol. Subsequently, samples were washed three times with PBS, and then blocked with PBS containing 10% FBS and 0.1% TRITON-X, 4°C overnight. The tissues were then incubated with primary mouse monoclonal anti-human cardiac troponin T (Thermo Fisher Scientific) or mouse monoclonal anti-human CD3 (Thermo Fisher Scientific), at 4 °C, overnight. The samples were then washed three times with PBS for 15 min, incubated with anti-mouse Alexa Fluor 488 (Life Technologies) and anti-mouse Alexa Fluor 647 (Abcam) secondary antibody, overnight, at 4°C, and then rinsed three times with PBS. For PBMC identification, it was tagged with CellTracker dye CM-DiI according to manufacture’s protocol. Immunofluorescence images were captured with a confocal fluorescence microscope (Nikon A1R).

### Cytokine Analysis

I3M medium was collected from the cardiac tissue compartment of the InVADE platform after viral infection. SARS-CoV-2 in the collected medium was inactivated by the addition of 1% Triton-X for 1 h, in accordance with the CL-3 facility safety protocol. Samples were then centrifuged at 500xg for 10 min and 30µl supernatant was collected for proinflammatory cytokine/chemokine analysis. The concentrations of IL-1α, IL-1β, IL-4, IL-6, IL-8, IL-10, IL-13, MCP-1, IFN-γ and TNF-α were measured using the Quantibody^®^ Human Inflammation Array (RayBiotech, USA), according to manufacturer’s protocol. The fluorescence was analyzed using GenePix^®^ Professional 4200 Microarray Scanner at 530nm.

### Functional Assessment of Cardiac Tissue

After 72 h of SARS-COV-2 infection, contractile activity of cardiac tissue was assessed by measuring activity in response to electrical field stimulation using an external stimulator (CStype 223, Hugo Sachs electronics-Havard apparatus). Excitation threshold (ET) was first measured, followed by measuring displacement of micro-cantilevers from videos recorder in the CL3 facility at 30 frames at 2xET at 1Hz. Relative changes in intracellular calcium concentrations were determined by adding Fluo-4 NW (Thermo Fisher Scientific) to the cardiac tissue compartment for 30 min, at 37 °C. Intracellular calcium transients were recorded at 30 frames at 2xET under green light channel (λ_ex_/ λ_em_= 490/525nm) at 1Hz. Videos are then analyzed using ImageJ Software (NIH). The calcium transient amplitude and kinetics of the calcium signals were analyzed using a customized MATLAB code as previously described^44^. The ratio of peak tissue fluorescence intensity, dF/F_0_ was calculated to determine the relative changes in intracellular Ca^2+^. The contractile measurements were determined using the ImageJ SpotTracker plugin. Fluorescent microscope (EVOS, Life technologies) was used to record videos in the CL- 3 facility.

### Study Participant and Data Collection

This study was approved by the Unity Health Toronto Research Ethics Board (REB no. 20-078). Forty patients with acute respiratory failure and suspicion of COVID-19 who admitted to medical-surgical intensity care units (ICUs) in Toronto, Canada between March 2020 and March 2022 and enrolled in COVID-19 Longitudinal Biomarkers of Lung Injury (COLOBILI) study. Infection status of admitted patients was confirmed prospectively by SARS-CoV-2 polymerase chain reaction tests and by antigen testing. The demographic characteristics (age and sex), cardiac examination (left ventricular ejection fraction, left ventricle size, right ventricle size, systolic function, and diastolic function) and clinical laboratory values (pH, pO_2_, and lactate) were recorded, along with the synchronous collection of blood samples. The cohort of forty patients was categorized into two different groups: SARS-CoV-2 negative patients requiring ICU care (n=20) and SARS-CoV-2 positive patients requiring ICU care (n=20).

### Circulating cell-free mitochondrial DNA Measurement

Mitochondrial DNA was extracted from 100µL cell supernatant and 50µL patient plasma samples following the manufacturer’s protocol provided with the QiaAMP DNA mini kit (Qiagen). Extracted DNA was eluted with ultra-pure distilled DNAse-free and RNAase-free water (Invitrogen): 50µL for cell supernatant and 100µL for patient plasma. A commercially synthesized oligonucleotide of the PCR product (Integrated DNA Technologies), of known concentration, was serially diluted to a concentration ranging from 10^8^ to 10^2^ copies/µL and used to estimate the absolute concentration of ccf-mtDNA. Mitochondrial genes, ND4 and ND1, were used to represent the major and minor arc of the mitochondrial genome. TaqMan^TM^ Duplex PCR was run on BioRad’s C1000 Thermal cycle CFX96 Real Time System using 20µL reaction mixture including 10µL TaqMan^TM^ Fast Advanced Master Mix (ThermoFisher), 4µL DNA, 1µL each of forward and reverse primers, and 1µL TaqMan^TM^ probe for each gene. qPCR cycling conditions were follows: 50 °C for 2 min, 95 °C for 20s, 40 cycles of 95 °C for 3s, and 60 °C for 30s, followed by a fluorescence read per cycle.

### Primer/Probe Sequence

ND4- F1 5’-CCATTCTCCTCCTATCCCTCAAC-3’

ND4- R1 5’-ACAATCTGATGTTTTGGTTAAACTATATTT-3’

ND4- Probe 5′-FAM/CCGACATCA/ZEN/TTACCGGGTTTTCCTCTTG/3IABkFQ/-3′ ND1- F1 5’-CCCTAAAACCCGCCACATCT-3’

ND1- R1 5’-GAGCGATGGTGAGAGCTAAGGT-3’

ND1- Probe 5′-HEX/CCATCACCC/ZEN/TCTACATCACCGCCC/3IABkFQ/-3′

ND4 + ND1 geneblock - 5’- CACGAGAAAACACCCTCATGTTCATACACCTATCCCCCATTCTCCTCCTATCCCTCAACCCCGACATCATTACCGGGTTTTCCTCTTGTAAATATAGTTTAACCAAAACATCAGATTGTGAATCTGACAACAGAGGCTCTCTTCACCAAAGAGCCCCTAAAACCCGCCACATCTACCATCACCCTCTACATCACCGCCCCGACCTTAGCTCTCACCATCGCTCTTCTACTATGAACCCCCCTCCCCATACCCAA-3’

### EV Isolation and Analysis

EVs were isolated using the miRCRY^TM^ exosome isolation kit (Qiagen), according to the manufacturer’s protocol. Briefly, 80% confluent HUVECs were washed once with PBS and switched to fresh serum-free culture medium, which was collected 48 h thereafter. The conditioned medium was centrifuged at 3200g for 10 min to remove cell debris. Supernatant was collected and 400µL exosome isolation reagent was added to 1ml media to allow EVs to precipitate overnight, at 4°C. The supernatant was discarded. Pellets were resuspended with PBS without mixing to wash off exosome isolation reagent, then centrifuged for 1 min at 3200g. The supernatant was then carefully aspirated. Precipitated EVs were reconstituted in PBS and the concentration and size distribution of 1x diluted EV samples were determined by nanoparticle tracking analysis (NTA) via NanoSight (Malvern). For all HUVEC-EV treatment studies, 2.5x10^8^ EVs were added to 1ml of cell culture medium.

### Western Blotting

Western blotting was performed using ExoA Antibody Kit according to the manufacturer’s protocol (System Biosciences, USA). Immunoblots were cut prior to primary antibody hybridization. Immunopositive bands were detected using the ECL Plus Kit (Invitrogen, USA), according to the manufacturer’s protocol. Antibodies were used at the following concentration: anti-CD9 (1:500; Cell Signaling), anti-CD63 (1:500; Abcam), anti-calnexin (1:500; Abcam), and anti-flotillin-1 (1:500; Cell Signaling). Images provided are the largest view saved from image acquisition.

### Transmission Electron Microscopy

EV pellets, isolated as described above, were suspended in 4% electron microscopy-grade paraformaldehyde in sodium phosphate buffer. EVs (10µL) were placed on square carbon electron microscopy grids for 20 min, after which excess solution was washed off with a filter paper. The sections were stained with saturated uranyl acetate for 5 min, and rinsed in distilled water, followed by Reynold’s lead citrate solution for 5 min. To image cardiac tissue sections, cardiac tissues were fixed with 10% formalin and 1% glutaraldehyde in 0.1M PBS for 1 h at room temperature. The samples were then fixed in 1% osmium tetraoxide in PBS for 1 h at room temperature, in the dark. Cardiac tissues were dehydrated using an ethanol series of 30%, 50%, 80%, and 95%, each for 5 min. The samples were then transferred into 100% ethanol three times for an additional 10 min, at room temperature, for complete dehydration. The samples were then washed twice with propylene oxide for 30 min, and then embedded in epoxy resin. After the polymerization, the solid resin block were sectioned on a Reichert Ultracut E-microtome to 90nm thickness and collected on 200 mesh copper grids. The sections were stained with saturated uranyl acetate and Reynold’s lead citrate solution for 15 min and then photographed with a Hitachi H7000 transmission electron microscope at an acceleration voltage of 80-120kV.

### EV miRNA sequencing

EVs were isolated from HUVEC-conditioned medium, as described above. EV pellets were lysed, and total RNA was isolated using the miRNeasy Micro Kit (Qiagen, cat# 217084), according to the manufacturer’s protocol. Total RNA was sent to Novogene for DNase treatment and subsequent preparation for sequencing. Briefly, treated total RNA was quantified using the Qubit RNA HS assay (ThermoFisher) and quality was measured using the Bioanalyzer 2100 Eukaryote Total RNA Nano assay (Agilent Technologies, CA, USA). Libraries were prepared using the QIAseq miRNA Library Kit (Qiagen, cat# 331502) and then sequenced using a NovaSeq S4 (Illumina) at a read length of 2x150bp and average depth of 12 million reads. R1 reads were trimmed to 75bp prior to analysis. Using the Qiagen RNA-seq Analysis Portal 3.0 (workflow version 1.2), trimmed R1 sequencing data were aligned to the human genome (GRCh38.103) via the miRbase v22 database. miRNA count files were extracted for further analyses.

### EV Labeling and Uptake by the Cells

EV suspensions were incubated with 2µM CellTracker dye CM-DiI (Invitrogen) in Hank’s Balanced Salt Solution (HBSS) for 5 min, at 37 °C, and subsequently placed on ice for an additional 15 min. Excess dye was removed using an exosome spin column (MW 100,000), as per the manufacture’s instructions. For uptake analysis, stained EVs from 0.5ml medium (approximately 0.5x10^9^ EVs) were cultured with HUVECs and PBMCs for 2 h. Cells were then rinsed twice with PBS, and then fixed with 4% paraformaldehyde. Immunofluorescence images were captured with a confocal fluorescence microscope (Nikon A1R).

### TLR Agonist Stimulation using THP-1

THP-1 dual-reporter cells (1x10^5^ cells/well in round bottom 96-well plates) were treated with a TLR agonist in combination with HUVEC-EVs: 1ng/ml diprovocim-1 (TLR1/2), 2µg/ml poly(I:C) (TLR3), 10ng/ml LPS (TLR4), 2µM ODN2395 (TLR9) and 1ng/ml TNF-α. For cytokine study, cytokines were added individually or in combination with HUVEC-EVs: 1ng/ml IL-6, 1ng/ml TNF-α and 1ng/ml IFN-γ. After a 24-h incubation, NF-kB and IRF activation were assessed by measuring the levels of SEAP and Lucia luciferase using Quanti-Blue and Quanti-Luc, respectively. SEAP levels were measured by reading optical density at 650nm. Lucia luciferase levels were determined by measuring the relative light units in a luminometer. HUVEC-EVs were added 2 hours prior to cytokine stimulation to determine the roles of EVs in modulating NF-kB and IRF activation.

### Measurement of extracellular ROS

1x10^5^ cells PBMCs were cultured in round bottom 96-well plate. PBMCs were then infected with SARS-CoV-2 at MOI of 1 for 24 h. To determine the role of EVs in modulating ROS generation, EVs were added 2 h prior to SARS-CoV-2 infection. Release of extracellular ROS was measured by plating cell culture supernatant in black 96-well plates, adding 2,7- dichlorofluoroscein acetate and incubating in room temperature. Fluorescence was read at 488nm excitation and 525nm emission.

### Proinflammatory Cytokine/Chemokine Stimulation of 3D Cardiac Tissues

Biowire cardiac tissues were prepared as previously described^44, 45^. Briefly, the Biowire platform consists of two parallel POMAC wires (100mm in diameter) placed at the opposing ends of ∼5mm long, 1mm wide microwell hot embossed into tissue culture polystyrene. To generate cylindrical cardiac tissue, human iPSC-derived cardiomyocytes were suspended in fibrinogen from human plasma (Sigma-Aldrich) at a concentration of 50x10^6^ cells/ml. Then, 1µL thrombin from human plasma was added to 3.5µL of the cardiomyocyte-human fibrinogen mixture (Sigma-Aldrich) and 2µL of the mixture is carefully pipetted into the microwell. The tissues were cultured for 7 days to allow for remodeling and tissue compaction in the presence of cardiac media described above. Subsequently, three tissue samples were placed in 12 well plates, each containing 1 mL of I3M media. Recombinant TNF-a, IFN-gamma, IL-6 and/or IL-8 (purchased from Peprotech) were added at concentration of 10ng/ml. 2.5x10^8^ EVs were co-treated with cytokine cocktails to determine the roles of EVs in modulating cardiac tissue functions. Cardiac tissue functions, including excitation threshold, maximum capture rate and contraction force were measured 6 h after the cytokine addition using Olympus CKX41 inverted microscope and CellSense Software. These measurements are performed under electrical pacing at 1Hz, via a pair of carbon electrodes (1/8 in in diameter, Ladd Research Industries) supplied by an external stimulator (Grass 88x) as we described in detail previously ^44^ . The autofluorescent properties of POMAC wires allow for determination of force of contraction by tracking displacement of the polymer wire at the center and conversion of displacement into force via calibration curves as we previously described. The displacement was measured from fluorescent videos captured at the 100 frames/s using what CMOS camera and Olympus CKX41 inverted microscope, and processed from individual frames using the ImageJ SpotTracker plugin. The force was calculated from displacement via calibration curves using a customized MATLAB code as previously described^44^.

### LDH Assay

The amount of lactate dehydrogenase (LDH) released by the cardiac tissues were analyzed with Cayman LDH cytotoxicity assay kit (Cayman), as per manufacturer’s protocol.

### Data and Statistical Methods

GraphPAD Prism version 9.0a was used for all statistical analyses and to generate plots. Data are reported as mean±SD. Significant differences between groups were analyzed by analysis of variance (ANOVA) with Bartlett’s test multiple comparison test and Student’s t-test, where appropriate. Differences at p<0.05 were considered statistically significant.

---

**Supplemental Figure 1:**
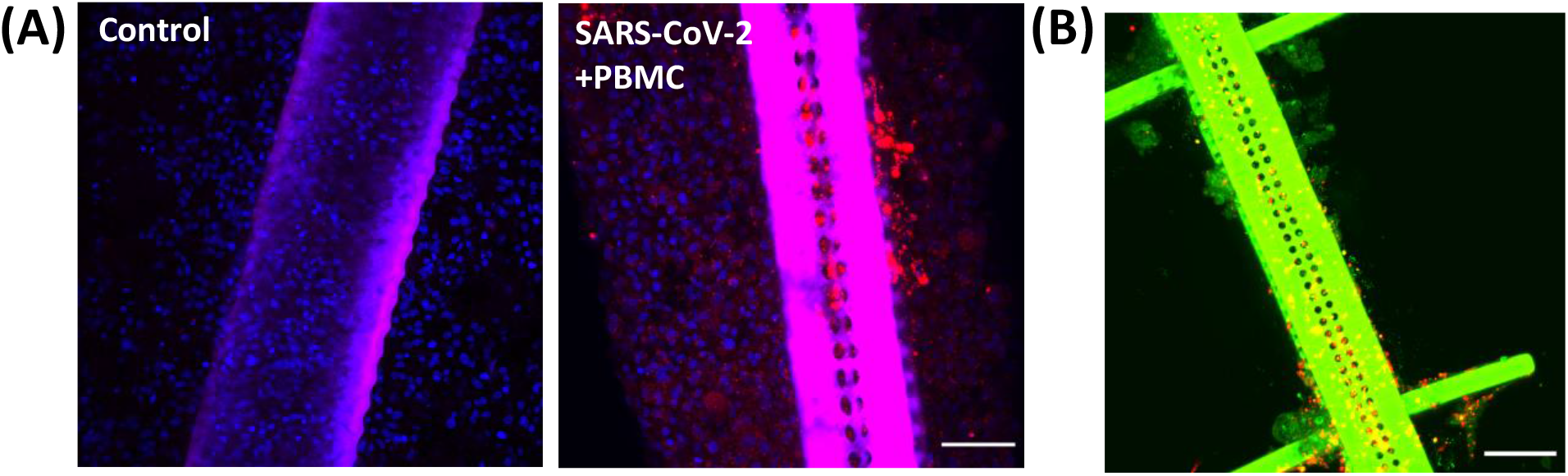
Immune cells infiltrate into engineered cardiac tissue. (A) Representative confocal fluorescence microscopy images of PBMC infiltrating the human iPSC-derived cardiac tissues upon SARS-CoV-2 infection. DiI-labelled PBMCs (red) and their DAPI-labeled nuclei (blue). Scale bar=100µm. (B) Representative confocal fluorescence microscopy images of infiltrated PBMC in the cardiac tissue. DiI-labelled (red) and CD3-labelled (green) PBMCs. Scale bar=200µm.

**Supplemental Figure 2:**
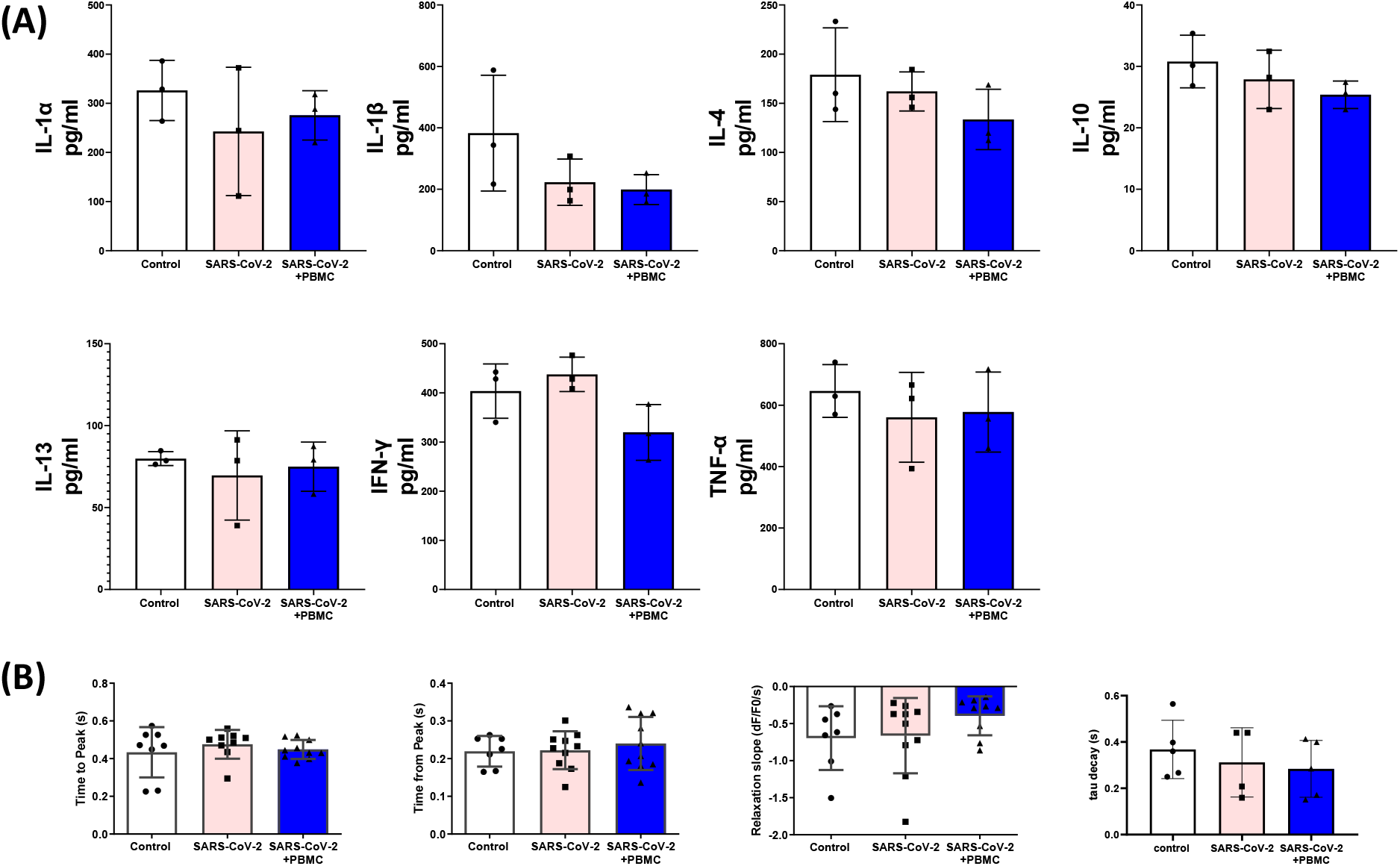
Analysis of proinflammatory mediators and intracellular Ca^2+^ kinetics in the inVADE heart tissues in the presence of SARS-CoV-2 and immune cells. (A) Measurement of cytokines and chemokines, and (B) intracellular Ca^2+^ kinetics 72 h after SARS-CoV-2 infection at MOI 0.1. Data are mean±SD, N=3. One-way analysis of variance (ANOVA) with Bartlett’s test, *p<0.05. No statistical difference was observed.

**Supplemental Figure 3:**
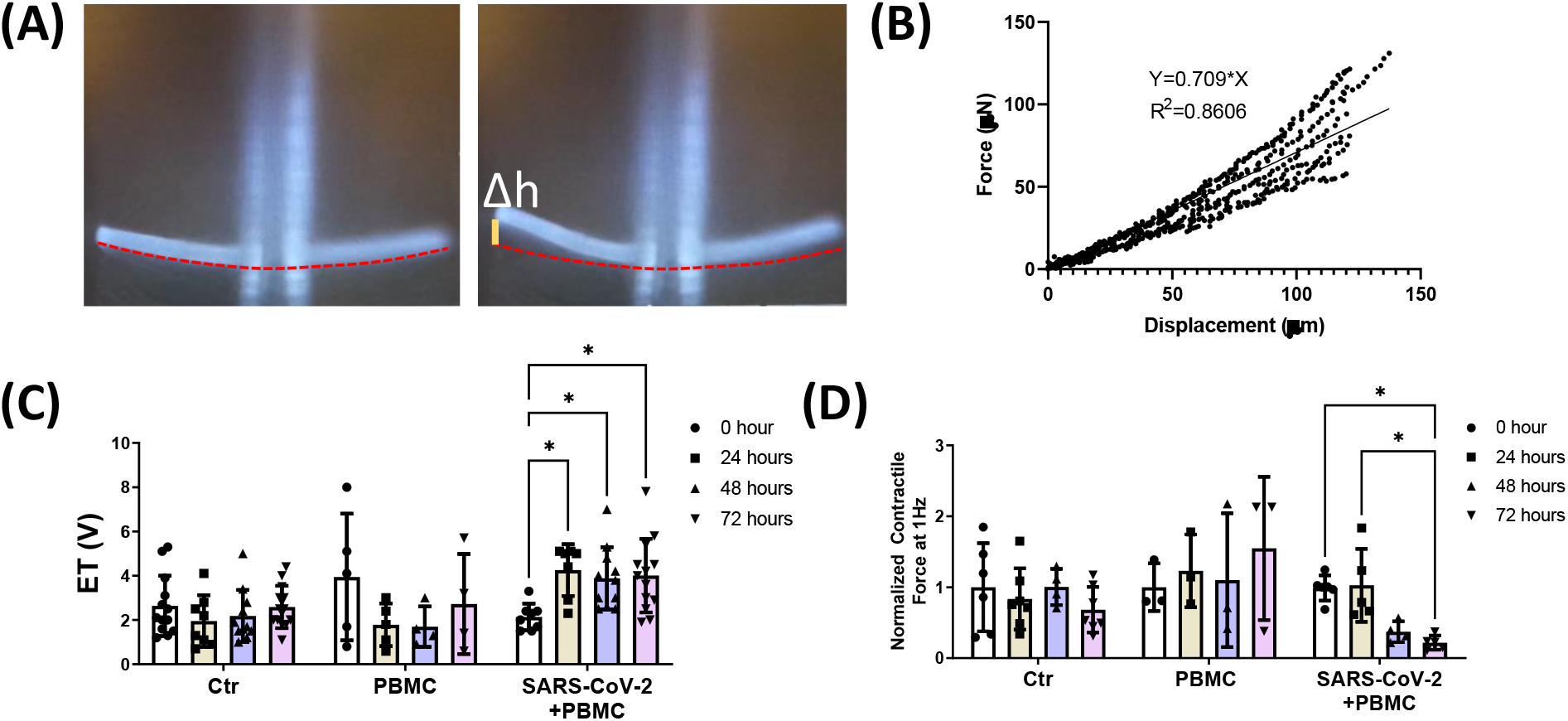
Analysis of contractile function in the InVADE cardiac tissues in the presence of SARS-CoV-2 and immune cells. (A) Representative images of autofluorescent POMaC cantilever structures observed under the blue channel during a contraction-relaxation cycle. Force generated by the cardiac tissue leads to displacement of cantilever (Δh) from the original position (red dashed line). (B) Representative force-displacement calibration curve of the cantilever structure as measured by a micro- mechanical tester. (C) Representative trace of cantilever displacement of the cardiac tissue being paced with increasing stimulation frequency 72 h after SARS-CoV-2 infection at MOI 0.1. (C) Excitation threshold and (D) normalized contraction force amplitude were measured under electrical stimulation at 0, 24, 48, and 72 h after SARS-CoV-2 2 infection at MOI 0.1. N=3. Two-way ANOVA with Bartlett’s test, *p<0.05, **p<0.01.

**Supplemental Figure 4:**
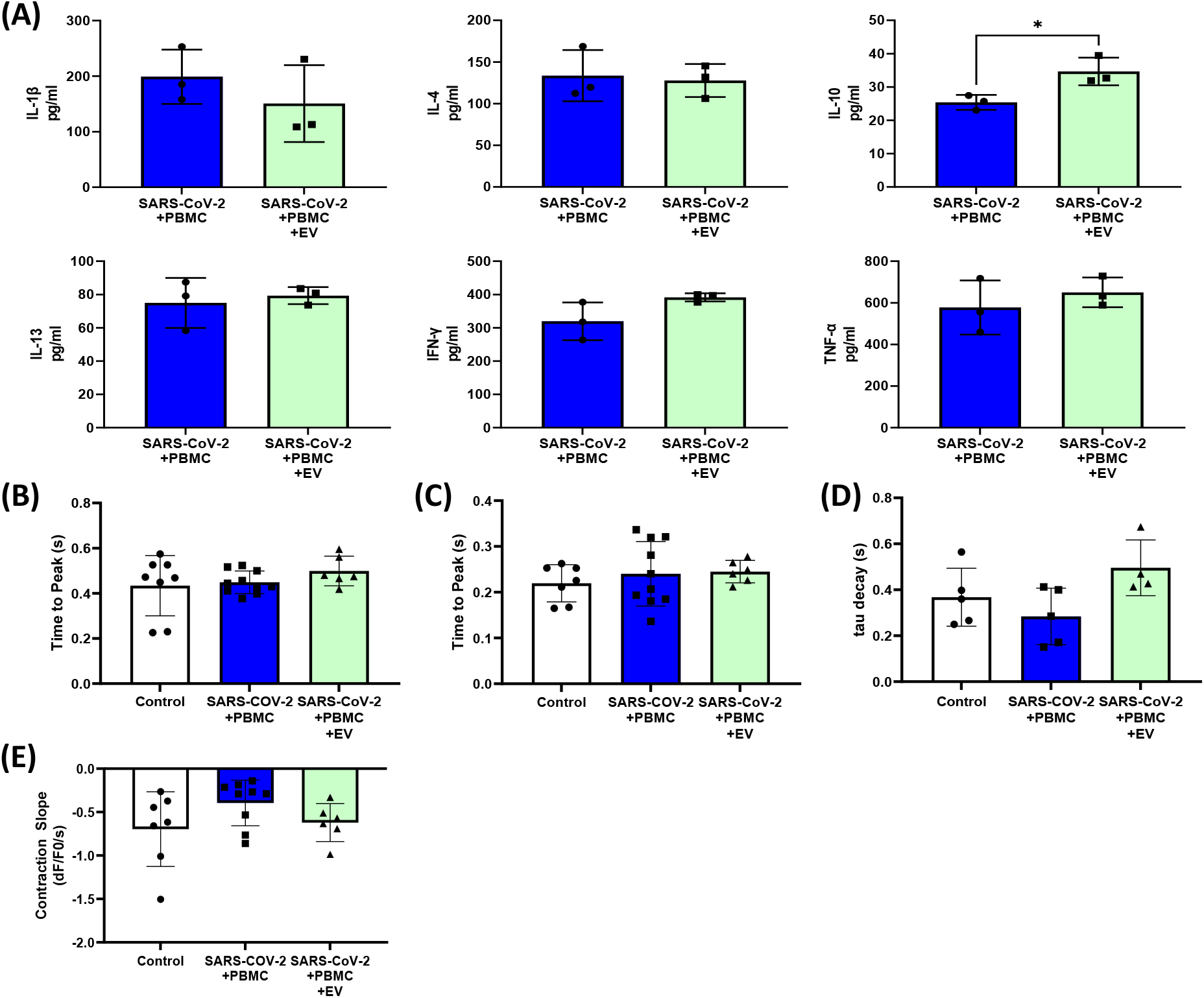
Analysis of inflammatory mediators and intracellular Ca^2+^ kinetics upon treatment of the SARS-CoV-2 infected tissues with exosomes in the presence of immune cells. (A) Measurement of cytokines and chemokines 72 h after SARS-CoV-2 infection at MOI 0.1 with HUVEC-EV treatment. Data are mean±SD, N=3. Student’s t-test with Bartlett’s test, *p<0.05. Analysis of intracellular Ca2+ kinetics 72 h after SARS-CoV-2 infection at MOI 0.1 with HUVEC-EV treatment: (B) time to peak, (C) time from peak, (D) tau decay and (E) relaxation slope. Data are mean±SD, N=3. One-way analysis of variance (ANOVA) with Bartlett’s test, *p<0.05.

**Supplemental Figure 5:**
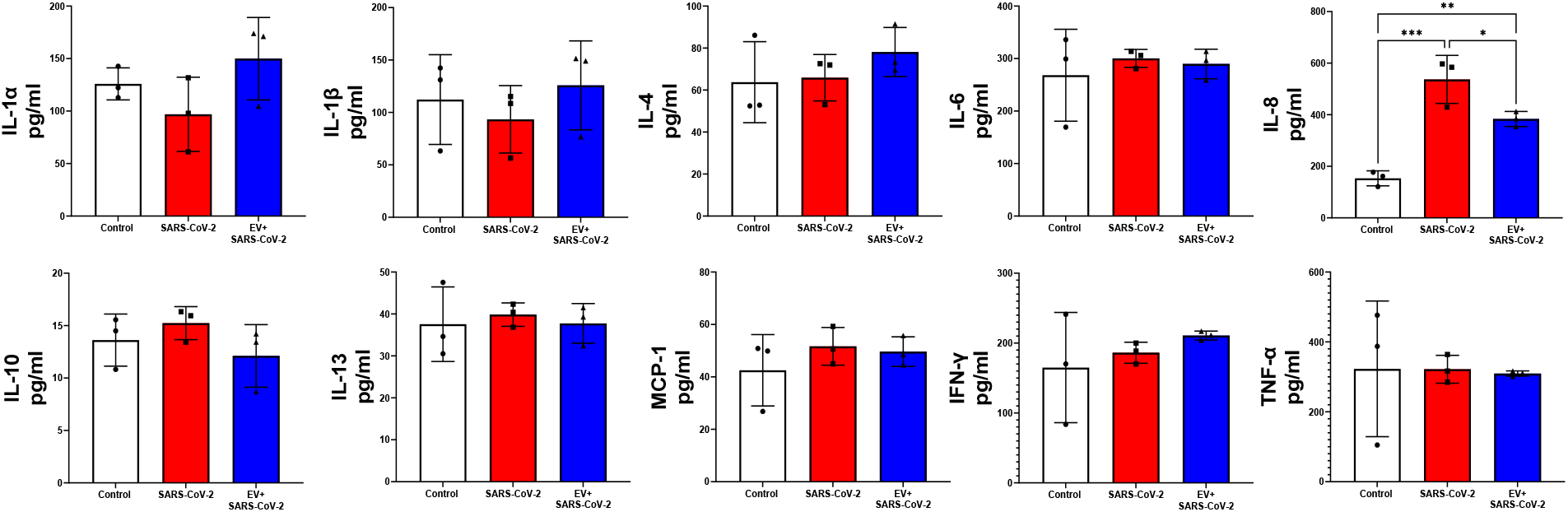
Analysis of proinflammatory cytokines and chemokines following SARS-CoV-2 infection of the cardiac tissues in the presence of immune cells. PBMCs were infected with SARS-CoV-2 at MOI of 1, and cell culture medium was collected 24 h after infection. Data are mean±SD, N=3. One-way ANOVA with Bartlett’s test, *p<0.05, **p<0.01, ***p<0.001.

**Supplemental Figure 6:**
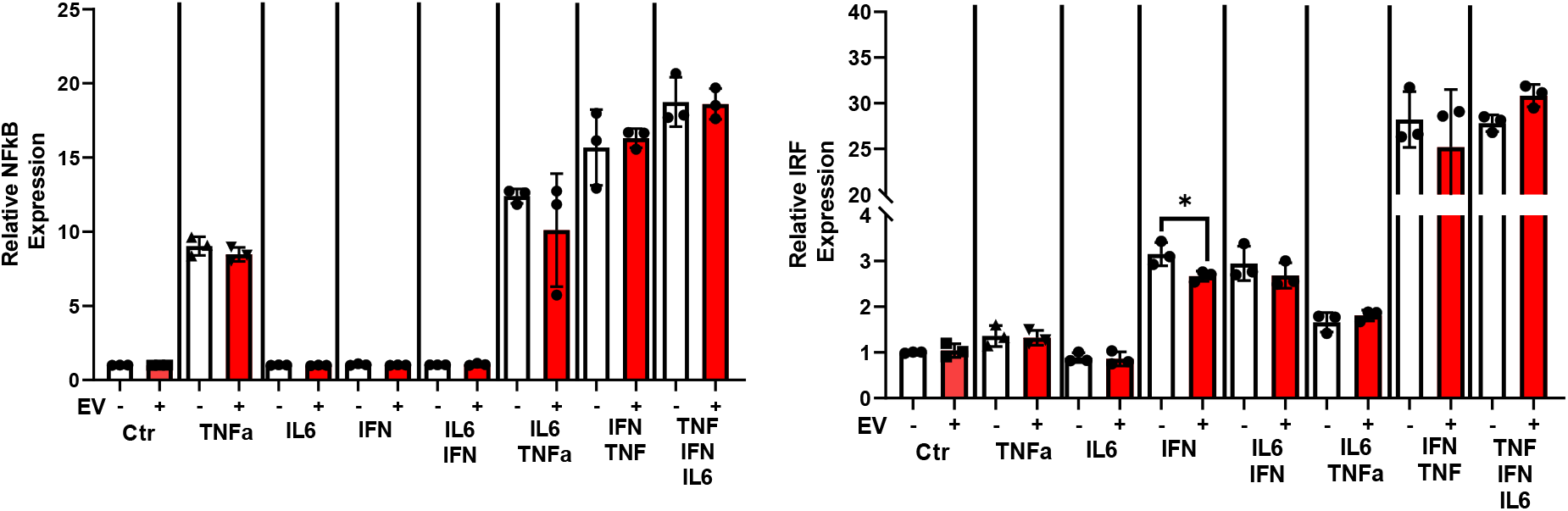
HUVEC-EVs do not repress NF-kB activation induced by TNF-α. THP-1 cells were first primed with EVs for 2 h. Cells were then stimulated with 1ng/ml IL-6, 1ng/ml IFN-γ and/or 1ng/ml TNF-α and incubated for 24 h. NF-kB and IRF activation were assessed by measuring the levels of SEAP and Lucia, respectively. Data are mean±SD, N=3. One-way ANOVA with Bartlett’s test, *p<0.05.

**Supplemental Figure 7:**
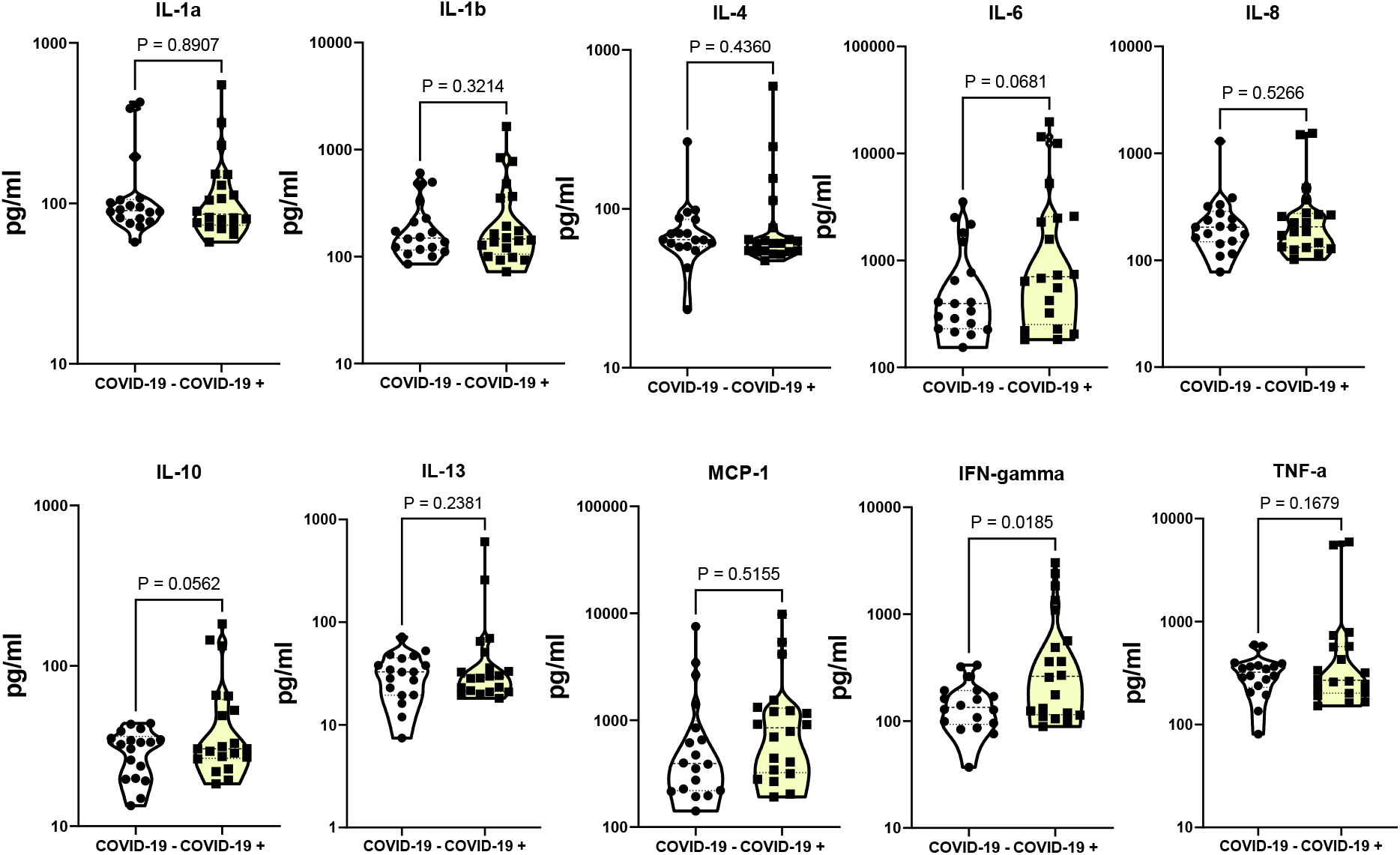
Quantification of cytokine and chemokine levels in COVID-19 patient plasma. Analysis of proinflammatory cytokines and chemokines in patient plasma samples. Data are mean±SD, n=20. Student’s t-test with Bartlett’s test, *p<0.05.

**Supplemental Figure 8:**
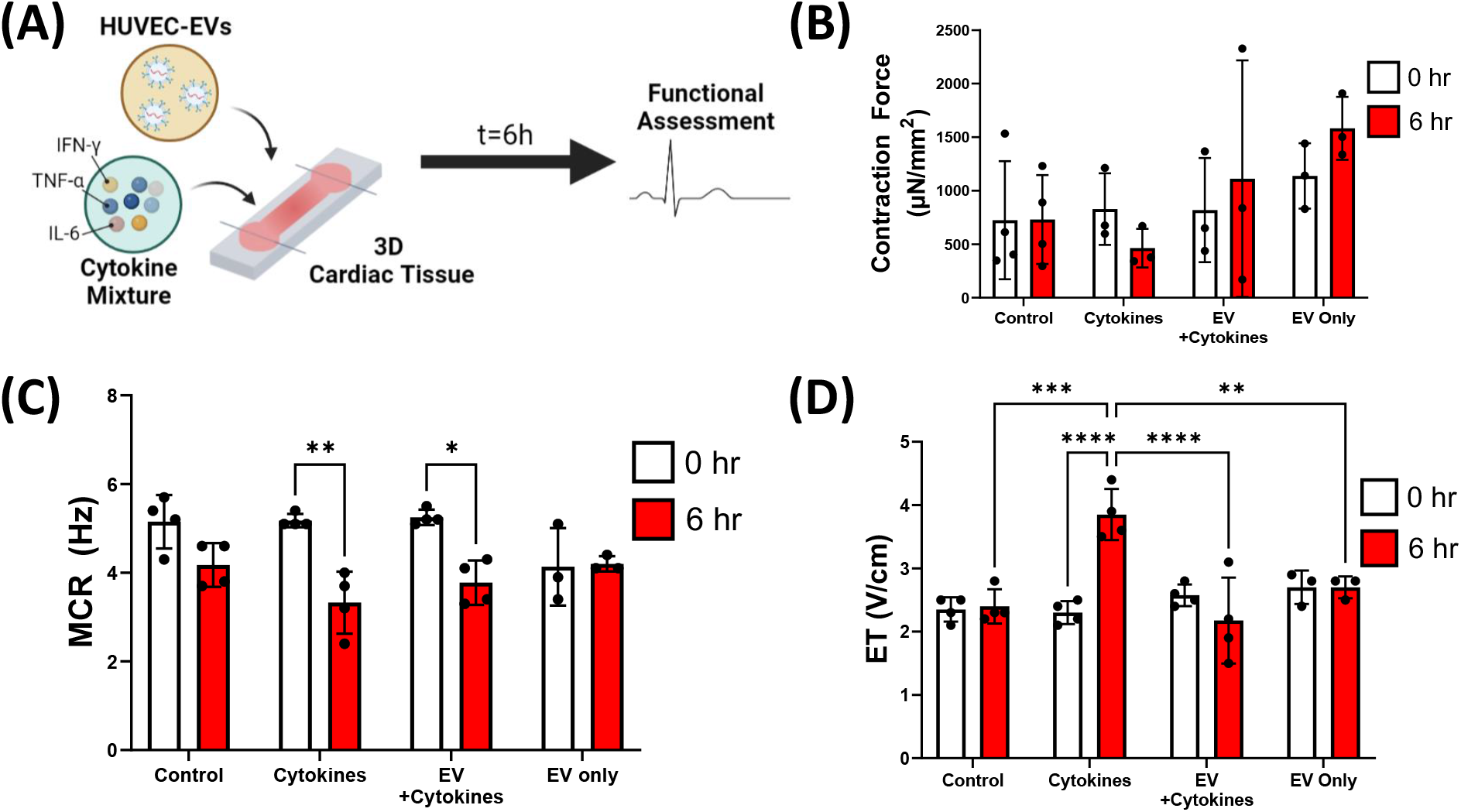
Direct HUVEC-EV treatment restores cytokine-mediated cardiac dysfunctions. (A) Schematic of the 3D cardiac tissues treated with cocktail of cytokines to simulate myocarditis. (B) Tissue contraction, (C) maximum capture rate and (D) excitation threshold were assessed before and after 6 h cytokine treatment. Data are mean±SD, n=3-4. Two-way ANOVA with Bartlett’s test, *p<0.05, **p<0.01, ***p<0.001, ****p<0.0001.

